# Exploring Thylakoid Emergence: Evolution of Membrane Biogenesis and Photosystem II assembly in early-diverging Cyanobacteria

**DOI:** 10.1101/2025.11.06.686923

**Authors:** Louise Hambücken, Denis Baurain, Luc Cornet

**Affiliations:** InBioS–PhytoSYSTEMS, Eukaryotic Phylogenomics, University of Liège, 4000 Liège, Belgium

**Keywords:** Photosynthesis, Thylakoids, Membrane Dynamics, Photosystem II Assembly, Cyanobacteria Evolutionary Biology, Phylogenomics

## Abstract

- Thylakoid membranes (TM) in cyanobacteria and chloroplasts host the light-dependent reactions of oxygenic photosynthesis. Gloeobacterales, the earliest-diverging cyanobacterial lineage, lack TM and perform photosynthesis in the cytoplasmic membrane, representing an ancestral state relative to other cyanobacteria (Phycobacteria). This study investigates the evolutionary origin of TM.
- Phylogenomic analyses were performed across a phylogenetically diverse set of cyanobacteria, including extensive representation of basal lineages (Gloeobacterales, Thermostichales, Gloeomargaritales and Pseudanabaenales), as well as micro- and macrocyanobacteria, using orthologous proteins involved in membrane dynamics and Photosystem II (PSII) assembly, together with structural modelling using AlphaFold3.
- We identified two candidate proteins associated with membrane trafficking that may contribute to TM biogenesis, including the SPFH family member Slr1106, proposed to have been acquired by lateral gene transfer. Analysis of 36 PSII assembly factors revealed modifications in late-stage assembly, notably in manganese homeostasis. Structural changes in the YidC translocase may have facilitated relocation of linear electron transfer components from the cytoplasmic membrane to TM.
- Altogether, these phylogenetic and functional prediction analyses provide new insight into the molecular innovations that led to TM emergence, including membrane trafficking systems, PSII assembly changes, and protein targeting adaptations.

## Introduction

Thylakoids are specialized membranes present in both cyanobacteria and chloroplasts that carry out the light-dependent reactions of oxygenic photosynthesis. They generate NADPH and ATP, for subsequent CO₂ fixation in the Calvin–Benson cycle, through a linear electron transport chain (LET) composed of multi-subunit complexes, notably Photosystem II (PSII) and Photosystem I (PSI), together with associated electron carriers. The emergence of this intracellular compartment provided cyanobacteria with an expanded membrane surface for light energy conversion, which is hypothesized to have contributed the Great Oxidation Event (GOE)^1^ approximately 2.4 billion years ago, profoundly transforming Earth’s biosphere^2^.

Among extant cyanobacteria, members of the order *Gloeobacterales,* the earliest-diverging lineage, lack thylakoid membranes (TM)^3,4^. Instead, they carry out oxygenic photosynthesis within specialized regions of their cytoplasmic membrane (CM)^5^. The absence of TM in this basal group suggests that these structures arose after the divergence of *Gloeobacterales* but before the emergence of *Thermostichales*, the second earliest-diverging cyanobacterial lineage, which already possesses TM. Consequently, *Gloeobacterales* retain the ancestral evolutionary state preceding TM formation and provide a unique model for exploring the transition from a LET located in the CM to a LET integrated within TM^6^. The proteins mediating the assembly of photosynthetic complexes within the TM may retain evolutionary signatures of this ancestral relocation, offering valuable insights into how cyanobacteria with TM, known as Phycobacteria, have emerged^6^.

The mechanisms underlying TM biogenesis remain poorly understood. Four main hypotheses have been proposed: fusion between CM and TM, vesicular transport, direct lipid transfer via soluble lipid carriers, or self-assembly driven by lipid phase transitions^1,7^ **(Figure 1)**. VIPP1, an homolog of the eukaryotic ESCRT-III system implicated in membrane deformation and scission, was suggested to play an essential role in the biogenesis of TM, thereby fitting the first scenario ^8–10^. This conclusion was supported by the fact that VIPP1 was only found in oxygenic photosynthetic organisms and that its presence is essential for thylakoid formation^10^. VIPP1 evolved from a duplication of the stress response protein PspA at the root of Phycobacteria and differs from PspA by an additional C-terminal extension^9^. However, a recent study showed that the *Gloeobacterales pspA* gene could complement a *vipp1-null* mutant in *Arabidopsis*^11^. Furthermore, proteomic data from TM biogenesis indicate that VIPP1 accumulates at late stages of TM formation^12^. Overall, these recent findings question the previously assumed central role of VIPP1 in the process.

**Figure 1.**
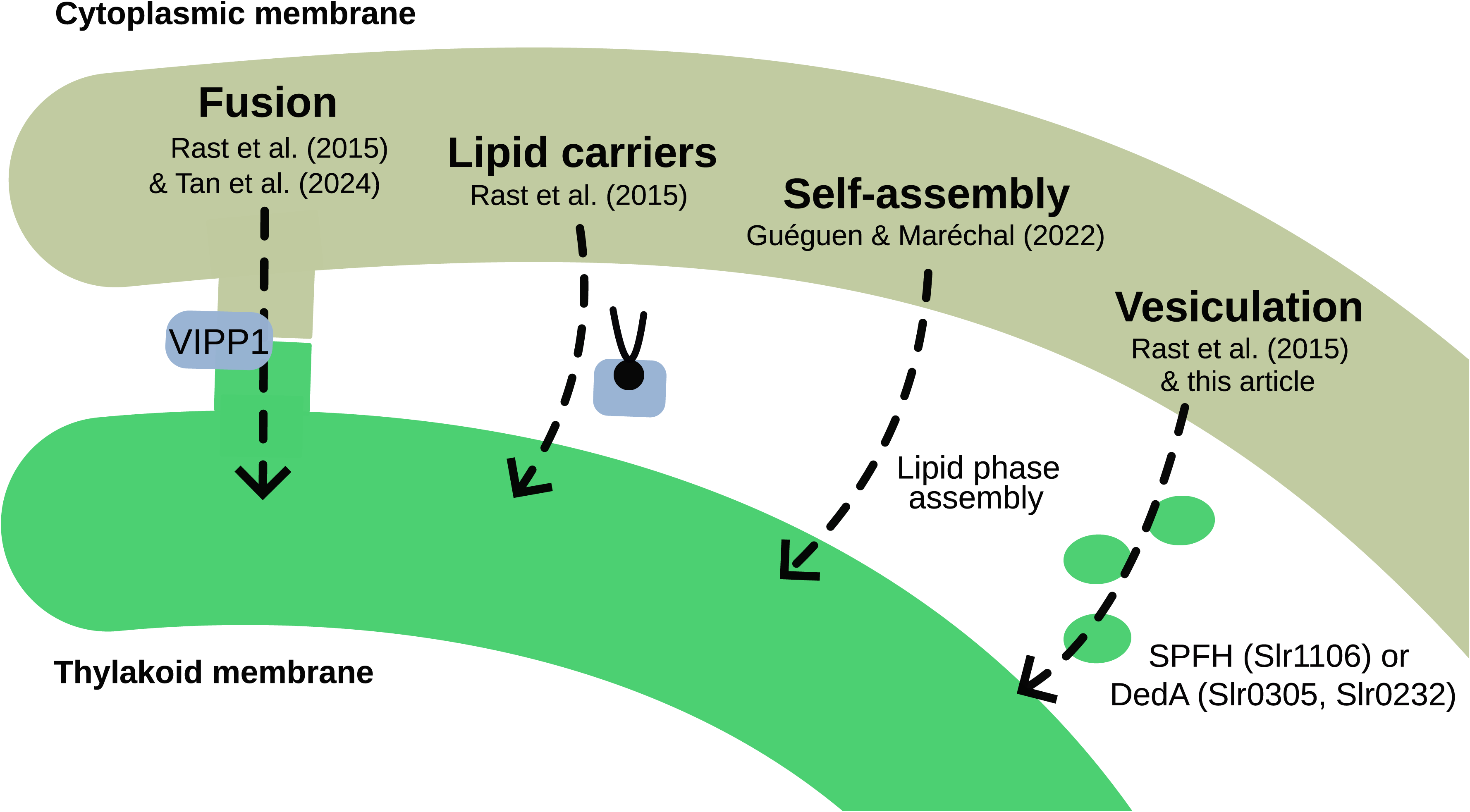
Proposed mechanisms of thylakoid membrane (TM) biogenesis. Schematic overview of the four main hypotheses proposed to explain TM biogenesis: (i) membrane fusion between the cytoplasmic membrane (CM) and the thylakoid membrane (TM) (7), (ii) direct lipid transfer mediated by soluble lipid carriers (7), (iii) self-assembly driven by lipid phase transitions (1), (iv) vesicular transport from the CM (7)

Although the mechanisms underlying the assembly of most of the protein complexes forming the LET remain poorly studied, *de novo* assembly of PSII was shown to be a highly synchronized spatial and temporal process in *Synechocystis*, mediated by at least 36 proteins known as assembly factors^13–15^. During the early steps of PSII biogenesis, the precursor of D1 (pD1) is translocated across the CM by the Sec translocon and the YidC/Alb3 insertase, followed by chlorophyll incorporation and D1 maturation. The D1–D2 heterodimer then forms within a specialized region of the CM enriched in assembly factors, known as the PratA-defined membrane (PDM)^16^. During the later steps of PSII biogenesis, the inner antenna subunits CP47 and CP43 are sequentially incorporated, and the Mn₄CaO₅ cluster is assembled and activated^17^. These later steps likely occur within the developing TM.

Here, we present a phylogenomic analysis of thirteen proteins involved in membrane trafficking that identifies one member of the SPFH family and one member of the DedA family as potential contributors to TM biogenesis, providing an alternative to the VIPP1-based model. We further report the first phylogenetic analysis of the 36 PSII assembly factors known so far, highlighting patterns of gene presence/absence and structural variations that suggest important modifications in the late stages of PSII assembly, notably in manganese homeostasis, after the divergence of *Gloeobacterales*. In parallel, we pinpoint structural changes in the translocase YidC and the Sec machinery that **may have** facilitated the relocation of the LET components from CM to TM.

## Materials and methods

### Genome selection and phylogenomic species tree

45,316 representative species genomes of Terrabacteria were retrieved from GTDB release r220 ^18^ (Figshare: https://doi.org/10.6084/m9.figshare.31135354). To limit computational burden during orthologous gene inference, a subset of 950 genomes were selected based on genome quality and phylogenetic position. Briefly, only the 43,827 genomes that passed the GUNC^19^ contamination filter were retained. Within each GTDB order, genomes were then ranked by the difference between their CheckM2^20^ completeness and contamination scores. For orders represented by a single genus, the two highest-quality genomes were kept; for all other orders, the best genome of each genus was selected, yielding 1,452 genomes in total, excluding the Cyanobacteriota phylum **(Table S1)**. Sampling within cyanobacteria was more exhaustive, resulting in 432 genomes, i.e., one or two representative genomes were chosen per genus, prioritizing those with >80% completeness and <10% contamination, as estimated by CheckM2^20^ **(Table S2)**. To ensure coverage of early-diverging cyanobacterial lineages, all available genomes from the orders *Gloeobacterales*, *Thermostichales* and *Gloeomargaritales* were incorporated, whereas the more abundant *Pseudanabaenales* were dereplicated using ToRQuEMaDA v0.2.1, with default parameters, resulting in 343 genomes^21^.

Protein-coding sequences were obtained directly from NCBI proteomes when available or predicted using Prodigal v2.6.3^22^ **(Table S3)**. A phylogenomic tree based on ribosomal proteins was reconstructed using the ORPER workflow with default parameters from the GEN-ERA toolbox v1 ^23^, with the exception that MAFFT v7.471 was used for the alignment of the ribosomal proteins^24^ and that the ribosomal tree was inferred using IQ-TREE v2.4.0^25^ with the best-fit substitution model (Q.YEAST+I+R10) and 1,000 ultrafast bootstrap replicates **(Figure S1)**. The resulting phylogeny was pruned using PARNAS v0.1.6^26^ to retain a final set of 950 representative species **(Table S4, Figures S2, S3)**, encompassing approximately 90% of the total phylogenetic diversity, as estimated by diversity score metrics **(Figure S4)**, and comprising 30 cyanobacterial species.

### Orthologous group inference

The orthologous groups were inferred with OrthoFinder v2.5.5^27^ from the 950 genomes selected using default parameters. An approach based on Hierarchical Orthologous Groups (HOGs) was first explored to account for multigene families stemming from duplications. However, this strategy, proved to be less accurate than the one based on standard Orthologous Groups (OGs), detailed below.

### Identification of the orthologous groups of interest

The OGs corresponding to known assembly factors and membrane dynamics proteins were identified by performing BLASTp searches (v2.9.0+)^28^ of every protein from the 950 proteomes against a database containing reference sequences of known PSII assembly factors and proteins involved in membrane dynamics (**Tables S5,6**). The e-value threshold was fixed at 10^-3^ and only the first hit was kept. The hits were then plotted based on their bitscore distributions for each OG **(Figures S5-S50)**. For each assembly factor and membrane dynamics protein, the OG with the highest median bitscore for cyanobacterial hits was selected **(Table S7)**. To check for optimal OG delineation, a reciprocal BLASTp of every reference sequence against a database containing the 950 proteomes was also performed. The best hits were then filtered based on their e-value, sequence length, number of copies per organism (first best hit per organism for the best score) and the fraction of the sequence covered by the alignment with the reference protein using OmpaPa (https://metacpan.org/dist/Bio-MUST-Apps-OmpaPa; D. Baurain). We then cross-validated the OGs derived from the filtered sequences against those obtained from the bitscore distribution plots.

### Orthologous gene tree inference

The gene trees of each OG of interest were generated using IQ-TREE v3.0.0^29,30^ with the best-fit substitution model and 1,000 ultrafast bootstrap replicates **(Table S7, Figures S51-S95)**. The sequences corresponding to the assembly factors and membrane dynamics proteins were then pinpointed in the gene trees based on the bitscores. When subsequent analyses were needed, either for taxonomic distribution determination or tree topology resolution, a HMM profile (pHMM) was built with the cyanobacterial sequences and searched against a database comprising the inferred proteomes of 343 Cyanobacteriota species (the dereplicated cyanobacterial dataset of 432 genomes, hereafter referred to as the cyanobacterial dataset) using HMMER v3.4^31^. The best hits were then filtered based on their e-value, sequence length, number of copies per organism (the three best hits per organism were kept) and the fraction of the sequence covered by the pHMM alignment (minimum 0.7) using OmpaPa. As the selected parameters varied depending on the protein under study, the filtering thresholds are detailed in figshare https://doi.org/10.6084/m9.figshare.31135354. Minimal lengths ranged from 8 to 556 amino acids, with a median of 154, and minimal –log10 e-values ranged from 0 to 281, with a median of 33.

The selected sequences were then aligned using MAFFT v7.505^24^. Alignment sites were conserved if they had at most 70% gaps and the sequences with a minimum of 30% length (with respect to the longest unaligned sequence) were exported using ali2phylip.pl (https://metacpan.org/release/Bio-MUST-Core; D. Baurain). Finally, a gene tree was inferred from the filtered alignment using IQ-TREE v3.0.0^29,30^ with the best-fit substitution model and 1,000 ultrafast bootstrap replicates **(Figures S96-S136)**. The presence or absence of the proteins was mapped onto a species tree of the cyanobacterial dataset^31^ using iTOL^32^.

### Functional protein and 3D protein structure prediction

Functional protein domains were annotated with InterPro v5.48-83.0^33^. When the experimental structures were not available, the 3D protein structures were modeled using the AlphaFold 3 server ^34^. The resulting structures were visualized and aligned using ChimeraX v1.11^35^. Electrostatic potential calculations were performed using the APBS plugin via the Poisson–Boltzmann server ^36^ (https://server.poissonboltzmann.org/) with the AMBER force field and default parameters.

All figures were created using Inkscape (Inkscape Project, n.d.).

## Results

### Membrane trafficking proteins

SPFH proteins (implicated in vesicular lipid transport), DedA family proteins (associated with lipid rearrangement), and dynamin-like proteins (involved in membrane fission and fusion) have been proposed by Siebenaller and Schneider (2023) as potential contributors to lipid trafficking in cyanobacteria^37^. To further explore their evolutionary history among Terrabacteria, we conducted a separate phylogenetic analysis for thirteen of these proteins **(Figure 2)**. Their taxonomic distribution revealed that two are absent from *Gloeobacterales* but present in most Phycobacteria included in this study: Slr1106 and Slr0305. The prohibitin Slr1106 is a member of the SPFH family. Its phylogeny within Terrabacteria (**Figure 3**) suggests for both proteins an origin through lateral gene transfer, possibly from Patescibacteria. Slr1106 appears to have been acquired at the base of the Phycobacteria, whereas Slr1768 may have been acquired later, prior to the emergence of the *Pseudanabaenales* but after that of the *Thermostichales* (**Figure 2**). Our analyses showed that the tree of Slr1106 and Slr1768 also contains two groups of basal Cyanobacteria, including *Gloeobacterales*, which lack orthologues in other cyanobacterial orders (**Figure 3**). Moreover, structural predictions revealed an additional β-sheet positioned outside of the ring structure formed during oligomerization in the proteins of the basal group I (**Figure S137**). The DedA protein Slr0305 was not detected in *Gloeobacterales* but was present from the root of Phycobacteria (**Figure 4**). Another DedA protein, Slr0232, is for its part conserved across all cyanobacteria. However, its phylogeny exhibits a long branch at the base of Phycobacteria (**Figure 5**), albeit without discernible structural or sequence divergence.

**Figure 2.**
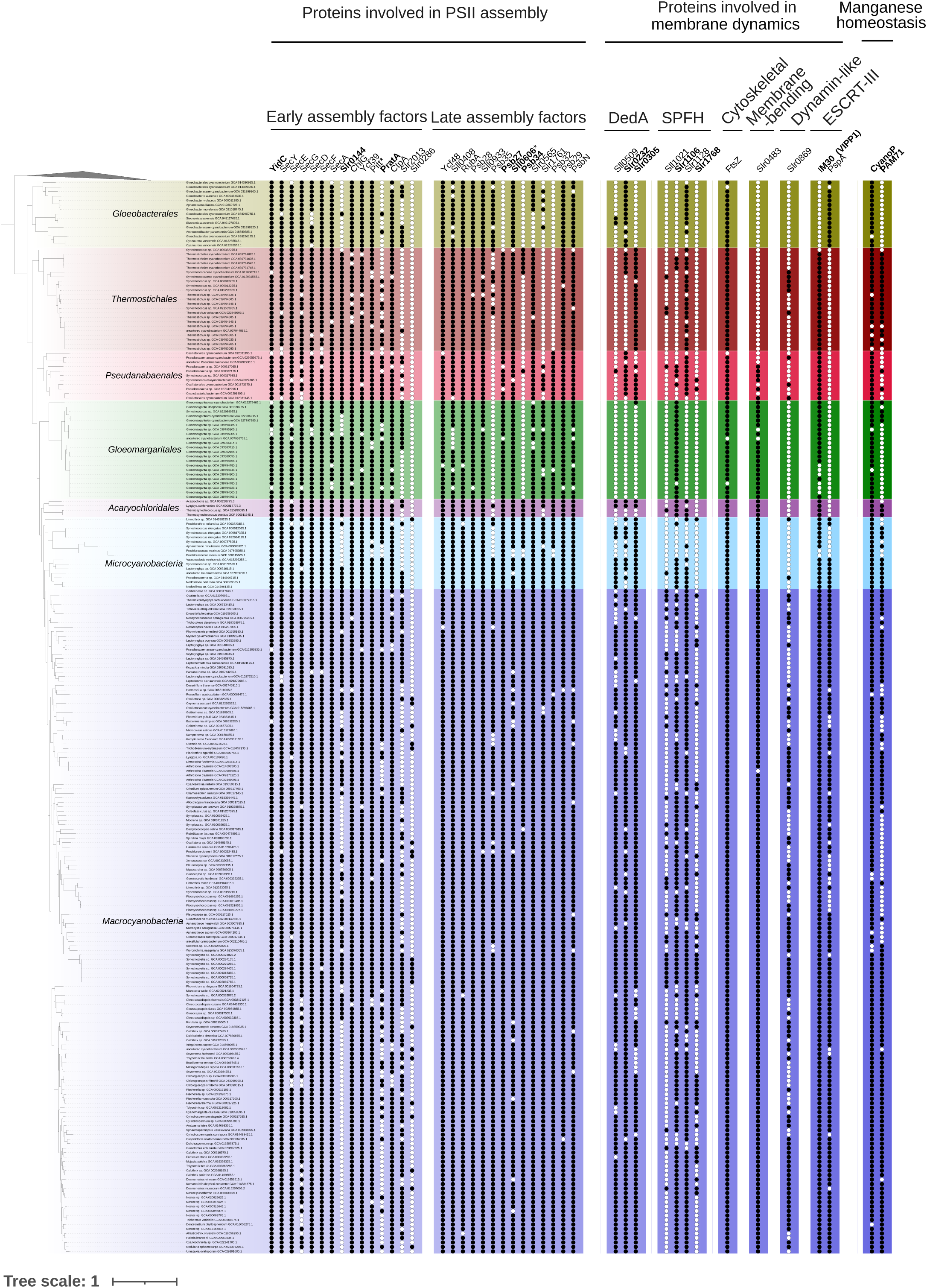
Taxonomic distribution of the PSII assembly factors and proteins involved in membrane dynamics in Synechocystis. A black dot indicates the presence of a protein, whereas a white dot indicates its absence. Highlighted proteins are discussed in the manuscript. Proteins marked with an asterisk indicate cases in which the Gloeobacterales homolog may not share the same evolutionary origin as the corresponding phycobacterial protein.

**Figure 3.**
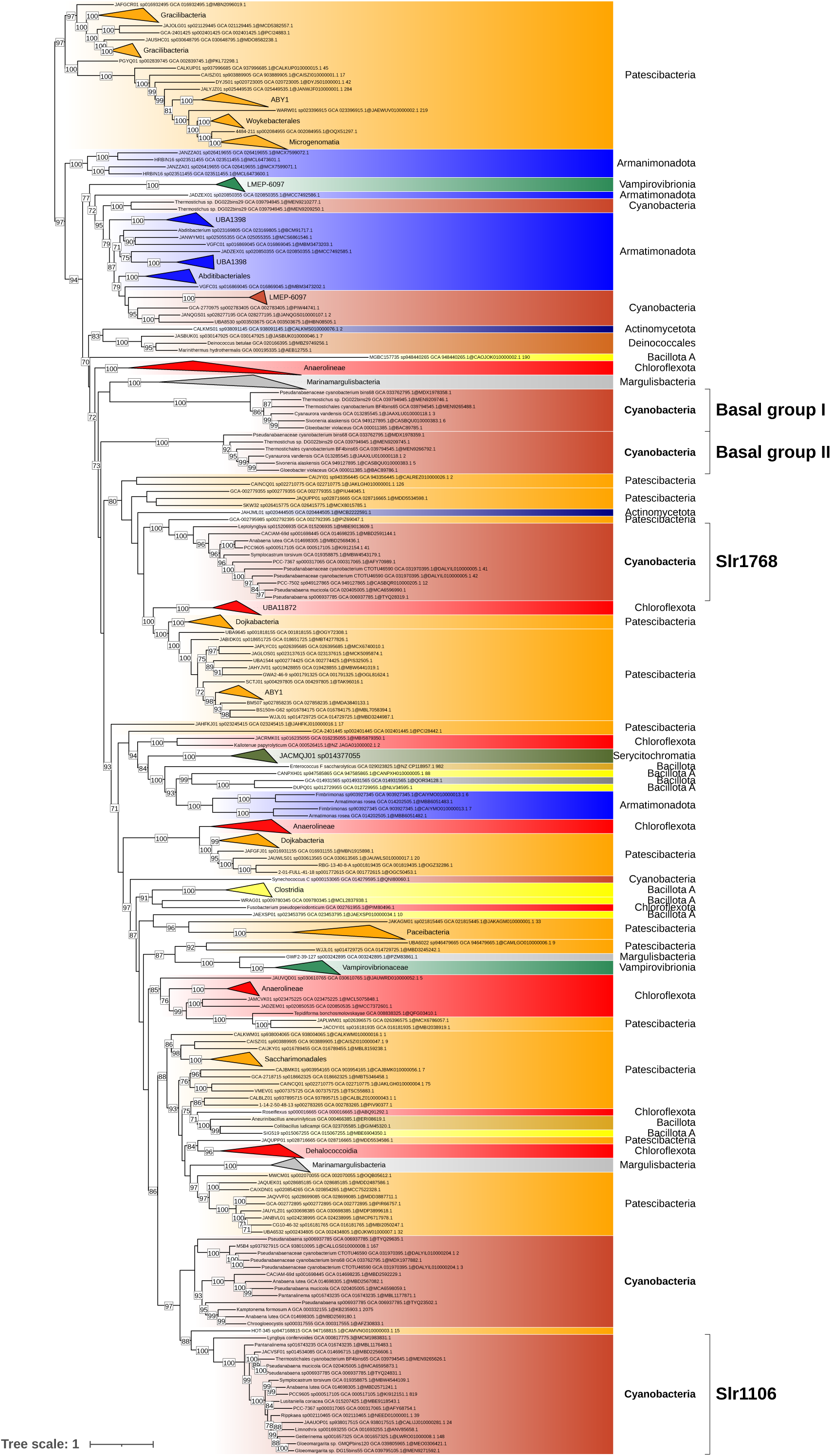
Phylogenetic tree of the cyanobacterial orthologous sequences of Slr1106 and Slr1768 (SPFH family) in the Terrabacteria dataset. The phylogenetic tree was constructed from 273 sequences across 510 unambiguously aligned positions using IQ-TREE (Q.PLANT+F+I+R6 model) with 1000 ultrafast bootstrap replicates. Bootstrap values ≥70% are shown. The tree annotation corresponds to the region showing the highest similarity to the reference Synechocystis sequences (NCBI Protein Accessions: BDH96643.1 [Slr1106] and BAA17070.1 [Slr1768]), as determined by BLAST.

**Figure 4.**
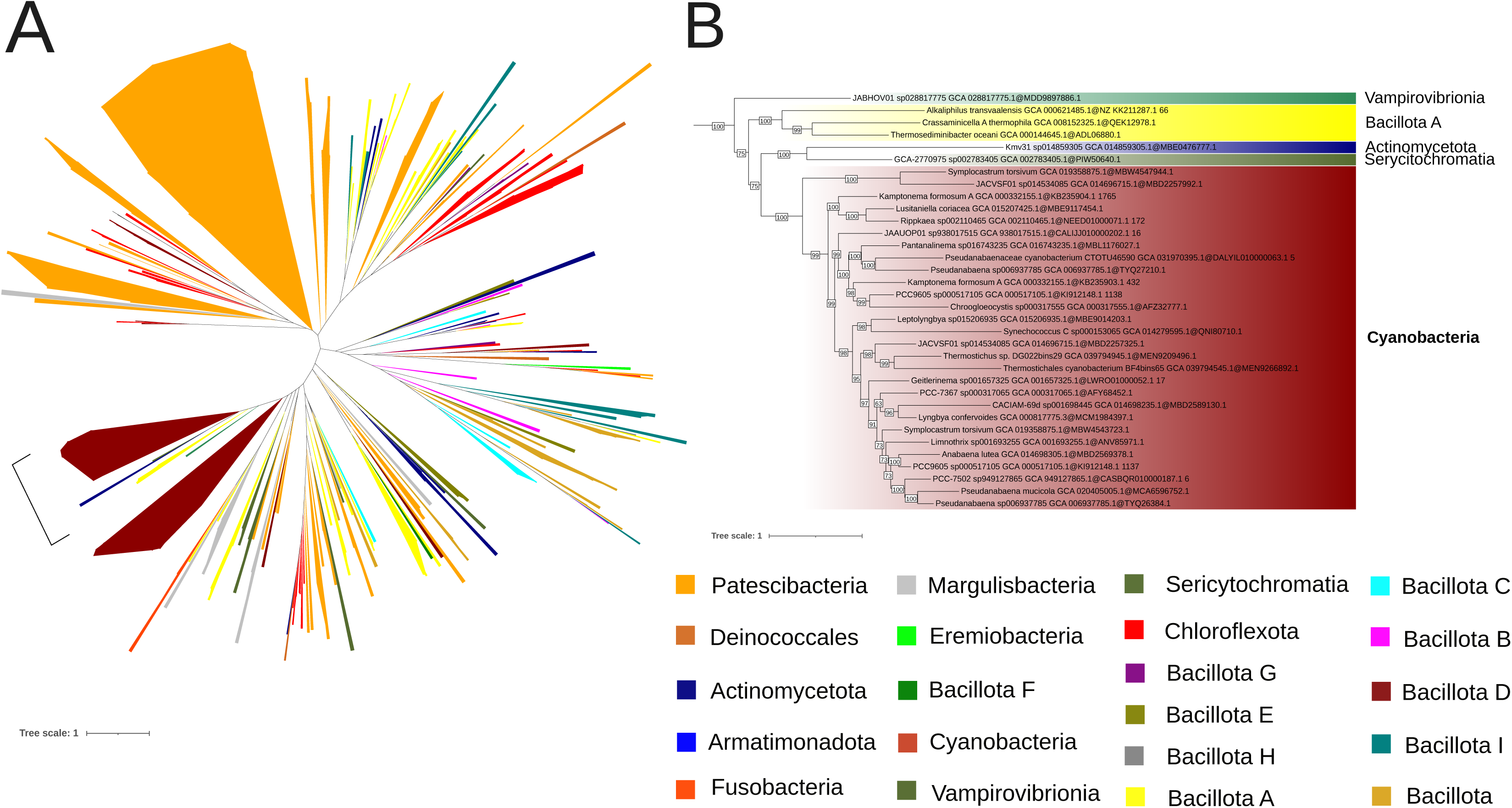
Phylogenetic tree of the cyanobacterial orthologous sequences of Slr0305 (DedA family) in the Terrabacteria dataset. (A) The phylogenetic tree was constructed from 347 sequences across 504 unambiguously aligned positions using IQ-TREE (LG+F+R7 model) with 1000 ultrafast bootstrap replicates. The region emphasized by brackets corresponds to the region showing the highest similarity to the reference Synechocystis sequence (NCBI Protein Accession: BAA10672.1) , as identified by BLAST. (B) The region labeled as Slr0305 in (A) was zoomed. Bootstrap values ≥70% are shown.

**Figure 5.**
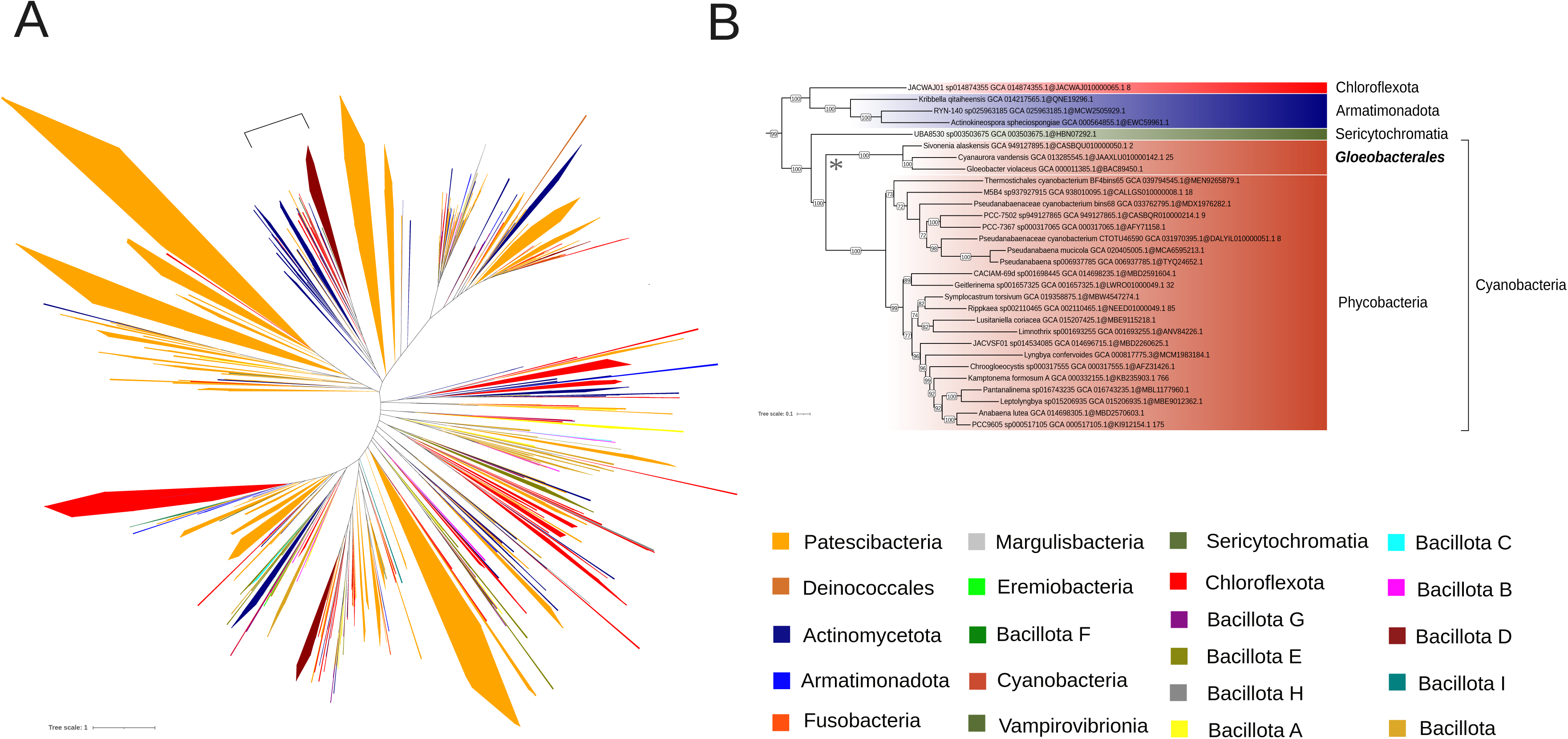
Phylogenetic tree of the cyanobacterial orthologous sequences of Slr0232 (DedA family) in the Terrabacteria dataset. (A) The phylogenetic tree was constructed from 987 sequences across 522 unambiguously aligned positions using IQ-TREE (LG+F+R10 model) with 1000 ultrafast bootstrap replicates. The region emphasized by brackets in the figure corresponds to the region showing the highest similarity to the reference Synechocystis sequence (NCBI Protein Accession: BAA10237.1), as identified by BLAST. (B) The region labeled as Slr0232 in (A) was zoomed. The long branch at the basis of Phycobacteria is indicated by an asterisk. Bootstrap values ≥70% are shown.

### Assembly factors of PSII

#### Late-stage assembly factors

The largest differences between *Gloeobacterales* and Phycobacteria were observed at the level of late-stage assembly factors **(Figure 2)**. Hence, Psb27 and Psb34 were not identified in *Gloeobacterales* in our analyses **(Figure 2)**. Psb34 is thought to bind CP47 and compete with HliA/B to enable the final stage of PSII assembly^38^, while Psb27 binds both unassembled and assembled CP43 in the PSII complex^39^. Sll0606, a protein of unknown function but leading to a massive reduction of D1 and CP43 in PSII assembly upon deletion^40^, was not detected either in *Gloeobacterales* (**Figure 2**). Domain analysis of Sll0606 suggests a potential role in the transport of hydrophobic components, and distant homologues in *Gloeobacterales* have been proposed^40^. In this work, we also identified such a homolog in *Gloeobacterales*: BAC90294.1. Its sequence is not orthologous to the *Synechocystis* PCC6803 reference sequence, but InterPro domain analysis annotates three domains capable of binding hydrophobic components at the N-terminal region **(Figure S138).** Hlips family phylogenies are not well resolved in our analyses, as evidenced by the lack of statistically supported clades separating HliA, HliB, HliC, and HliD. Instead, these proteins were scattered over several OGs **(Figure S9)**, and a combined phylogenetic tree did not reveal clear distinct monophyletic groups for each gene **(Figure S55)**.

Except for the accession GCA_038245785.1 (*Gloeobacterales* cyanobacterium), *Gloeobacterales* were also shown to lack Slr0144, a protein containing a 4-vinyl reductase (V4R) domain that binds small hydrophobic molecules and is inferred to interact with chlorophyll precursors in *Rhodobacter capsulatus*^41,42^. In *Synechocystis* sp. PCC6803, Slr0144 is encoded within the *slr0144*–*slr0152* operon^41^, which is involved in PSII stabilization^43^. Slr0144 and Slr1471, the only two proteins of *Synechocystis* sp. PCC6803 containing a V4R domain, are the result of a gene duplication (**Figure 6**). Slr0144 and Slr1471 differ notably by Slr0144 having an N-terminal extension, along with changes in its C-terminal region (**Figure S139**).

**Figure 6.**
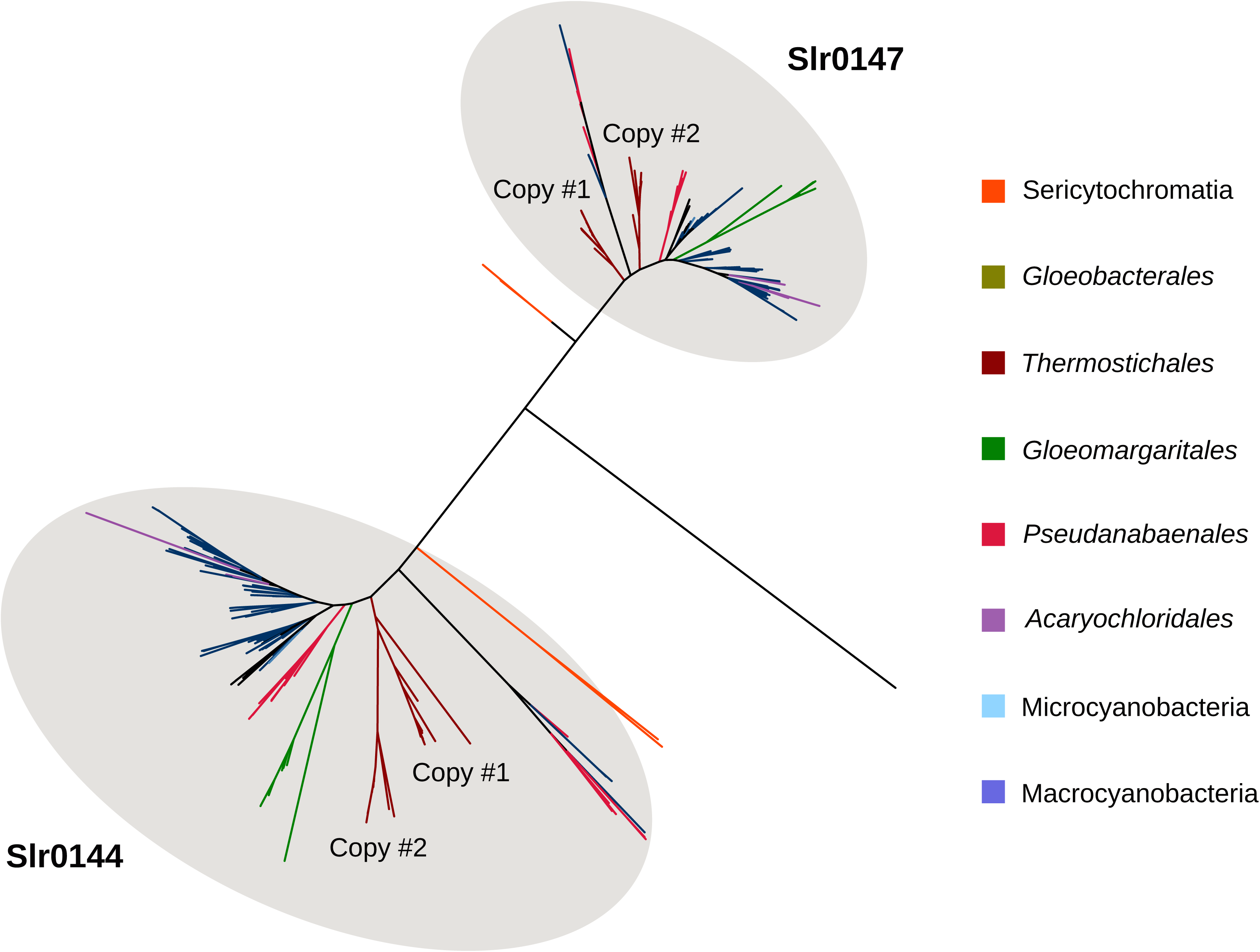
Phylogenetic tree of the cyanobacterial orthologues sequences of Slr0144 and Slr0147 in the cyanobacterial dataset. The phylogenetic tree was constructed from 311 sequences across 234 unambiguously aligned positions using IQ-TREE (Q.PFAM+R6) with 1000 ultrafast bootstrap replicates. The tree annotation corresponds to the region showing the highest similarity to the reference Synechocystis sequences (NCBI Protein Accessions: BAA18544.1 [Slr0144] and BAA18547.1 [Slr0147]), as determined by BLAST.

#### Early-stage assembly factors

The early stages of PSII biogenesis do not show any differences in terms of presence or absence of assembly factors between *Gloeobacterales* and Phycobacteria. Indeed, whether considering the processing and incorporation of the D1 precursor (pD1), D2 folding, or the formation of the reaction center, all corresponding assembly factors are present in *Gloeobacterales* (**Figure 2**). However, a structural difference exists in the periplasmic domain of the translocase YidC and SecD between *Gloeobacterales* and Phycobacteria, with the latter harboring a specific features in its periplasmic domain.

YidC is part of the Oxa1/Alb3/YidC family of proteins, which are involved in the insertion and assembly of membrane proteins and have homologs across all three domains of life ^44^. Notably, bacterial YidC shares homology with mitochondrial inner membrane Oxa1 and chloroplast thylakoid Alb3 proteins. YidC has been shown to participate in PSII biogenesis alongside the SecYEG machinery by mediating the membrane integration of the D1 precursor (pD1) in *Synechocystis* PCC6803, a process homologous to the one described in chloroplasts^45^. In both cyanobacteria and chloroplasts, this process involves the SecYEG translocon, YidC, SecDF, and SecA. SecYEG forms the core membrane channel for protein translocation; YidC acts as a membrane insertase facilitating the integration of membrane proteins; SecDF is an accessory complex that enhances translocation using the proton motive force; and SecA is an ATPase that drives preprotein translocation through SecYEG^46^. Notably, both YidC and SecDF contain periplasmic domains. The periplasmic domain of YidC is only present in diderm bacteria, such as cyanobacteria, as its size is markedly reduced in other bacterial and chloroplastic homologs^44^. Interestingly, our results indicate that the periplasmic domain of cyanobacterial YidC is largely conserved across the cyanobacterial phylum, except in *Gloeobacterales.* Sequence alignment reveals well-conserved N-terminal and C-terminal regions; however, the periplasmic domain of *Gloeobacterales* YidC differs, with missing amino acids in the sequence (**Figure S140**). Besides, 3D structure prediction of the periplasmic domain highlights these differences, showing that Phycobacteria YidC possesses a P1 domain with structural changes and two additional helix loops compared to the *Gloeobacterales* homolog, one predicted with confidence (pLDDT ≥ 70) and the other with low support (70 > pLDDT ≥ 50) **(Figure 7)**. In order to investigate how these changes affect YidC interaction with the Sec machinery, we performed the same analyses for SecD and SecF, the only two components of the Sec machinery that harbor periplasmic domains^47^. Our data show that SecD is predicted to contain an additional helix–loop insertion in *Thermostichales*, supported with relatively low confidence (70 > pLDDT ≥ 50). with this loop being even larger in *Synechocystis* and modeled with high confidence (overall pLDDT ≥ 70). **(Figure S141)**. In contrast, no structural differences were observed in the periplasmic domain of *Thermostichales* SecF, whereas such differences were identified in *Synechocystis* **(Figure S142)**.

**Figure 7.**
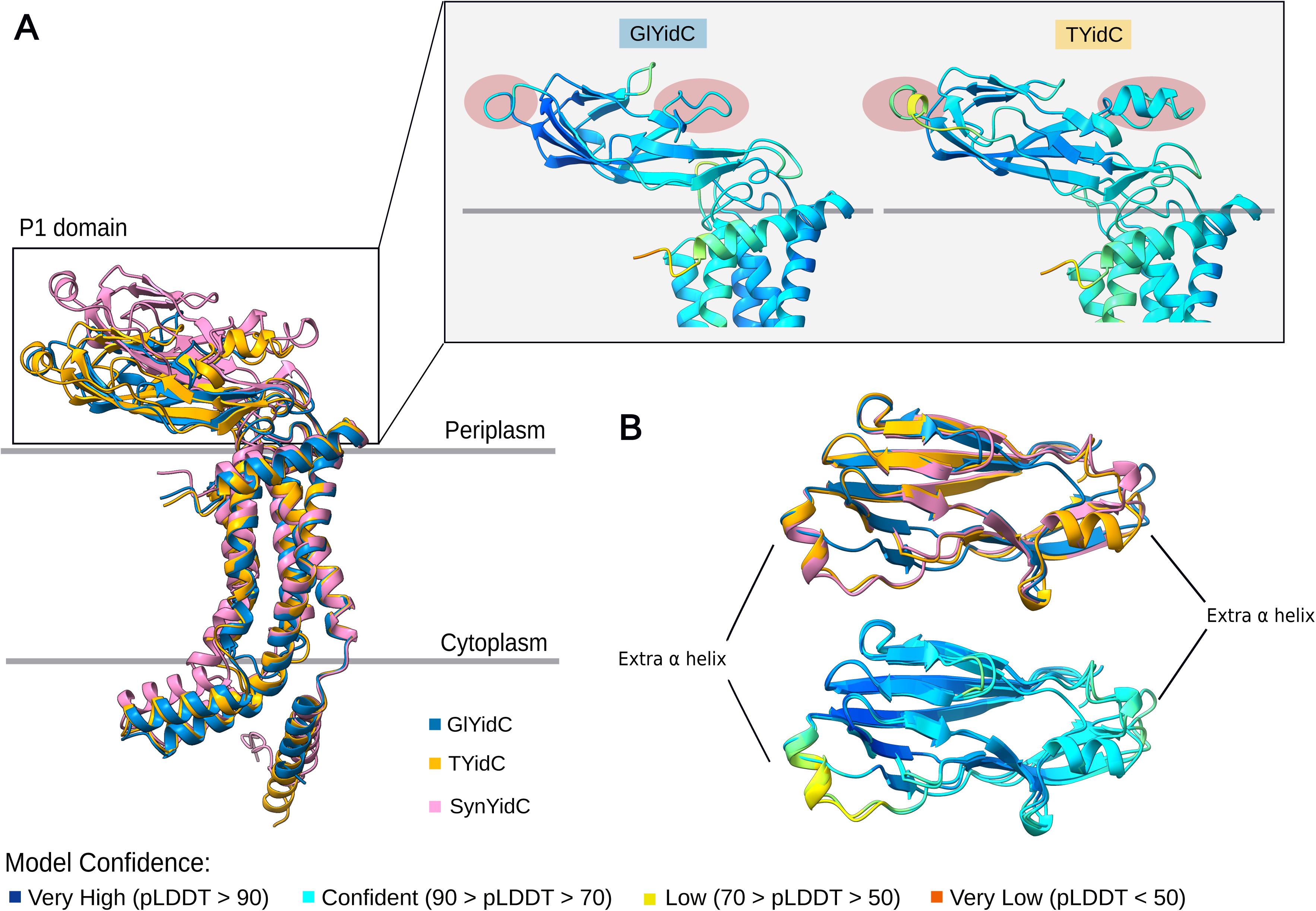
AlphaFold Prediction of the 3D structures of Gloeobacter violaceus PCC7421 (BAC90172.1–GlYidC), Thermostichales cyanobacterium GCA_039794945.1 (MEN9209976.1–TYidC) and Synechocystis sp. PCC 6803 substr. Kazusa (P74155.1 SynYidC). (A) Structure alignment of GlYidC (blue), TYidC (yellow) and SynYidC (pink). Positions of the additional α helices are highlighted in red. (B) Top view of the structure alignment of GlYidC, TYidC and SynYidC periplasmic domains. The Predicted Local Distance Difference Test (pLDDT) scores mapped onto the GlYidC and TYidC indicate confidence levels of the predicted models.

PratA, a periplasmic tetratricopeptide repeat (TPR) protein strongly associated with the PDM region^17^, has been shown to facilitate efficient Mn delivery to pD1^48^. Our taxonomic distribution analyses showed the absence of PratA in most *Gloeobacterales,* at the exception of two accession: GCA_018389385.1 (*Anthocerotibacter panamensis*) and GCA_038245785.1 (*Gloeobacterales* cyanobacterium) **(Figure 2)**. CyanoP, the cyanobacterial homolog of the PsbP protein known in land plants, is a Mn-binding protein that plays a role either in the binding association of CP47 and CP43 or in the photoactivation of the oxygen-evolving complex^49^. While our analyses show the presence of CyanoP in *Gloeobacterales*, its sequence has diverged substantially compared to Phycobacteria (**Figure S143**). APBS electrostatics calculations suggest that *Gloeobacterales* CyanoP either binds divalent ions differently compared to its Phycobacteria counterpart or does not bind ions at all (**Figure 8**). PAM71 is a Mn transporter involved in manganese homeostasis, sequestering excess cytoplasmic manganese into the periplasmic and lumenal compartments^50^. Its role in Mn uptake is considered to support PSII oxygen-evolving complex performance, suggesting a PratA-independent Mn delivery pathway^51^. Our results indicate its absence in *Gloeobacterales* **(Figure 2)**. MncA plays an important role in manganese binding within the periplasmic space^52^. However, its OG could not be determined, as we could not extract an OG that contained most cyanobacterial sequences with high sequence similarity **(Figure S9)**.

**Figure 8.**
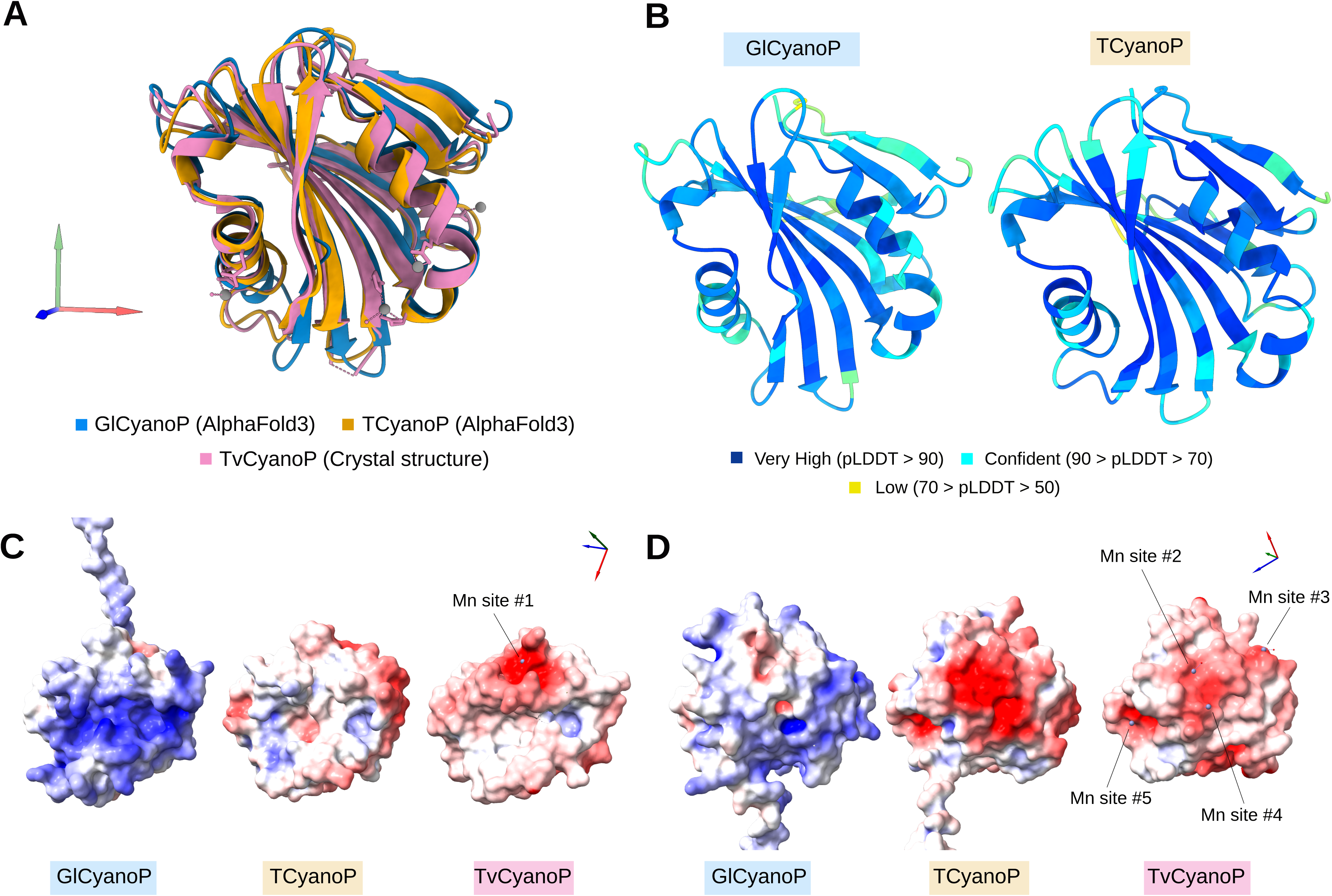
AlphaFold Prediction of the 3D structures of Gloeobacter violaceus PCC7421 (BAC89381.1–GlCyanoP), Thermostichales cyanobacterium GCA_039794945.1 (MEN9209059.1–TCyanoP) and PDB 2XB3 CyanoP structure from Thermosynechococcus vestitus BP-1 (TvCyanoP) with APBS electrostatics estimation. (A) Structure alignment of the predicted GlCyanoP (blue) and TCyanoP (yellow) AlphaFold structures and the TvCyanoP crystal structure (PDB: 2XB3) (pink). (B) Predicted Local Distance Difference Test (pLDDT) scores mapped onto CyanoP structures GlCyanoP and TCyanoP, indicating confidence levels of the predicted models. (C, D) APBS electrostatics estimation of GlCyanoP, TCyanoP and TvCyanoP structures with two different views displaying the putative Manganese-binding sites. The GlCyanoP and TCyanoP predicted structures were aligned to TvCyanoP in order to find the position of the ion-binding sites. The APBS electrostatics was estimated based on the APBS plugin via the Poisson–Boltzmann server (https://server.poissonboltzmann.org/) with the AMBER force field and default parameters.

## Discussion

To facilitate interpretation, the key results of this study are summarized in **Figure 9**, which outlines a possible evolutionary scenario. The emergence of TM is necessarily linked to a biological innovation involving lipid environments^1,7^. VIPP1 has long been considered a key factor in TM formation, a hypothesis supported by Tan *et al.* (2024), who demonstrated that VIPP1 originated from a duplication and subsequent C-terminal extension of the stress response protein PspA at the root of Phycobacteria^9^. However, the functional role of VIPP1 remains ambiguous. In the two model cyanobacteria *Synechocystis* sp. PCC 6803 and *Synechococcus* sp. PCC 7002, VIPP1 does not appear to participate in TM biogenesis but rather in the assembly of photosynthetic complexes^53,54^. This observation is consistent with the proteomic analysis of TM by Huang *et al.* (2023), which detected VIPP1 only during late stages of TM development^12^. Moreover, a recent study identified a PspA homolog in *Gloeobacterales* capable of complementing a *vipp1-null* mutant in *Arabidopsis*^11^. Although its function is still undefined, these results collectively indicate that VIPP1 is unlikely to be directly involved in TM formation.

**Figure 9.**
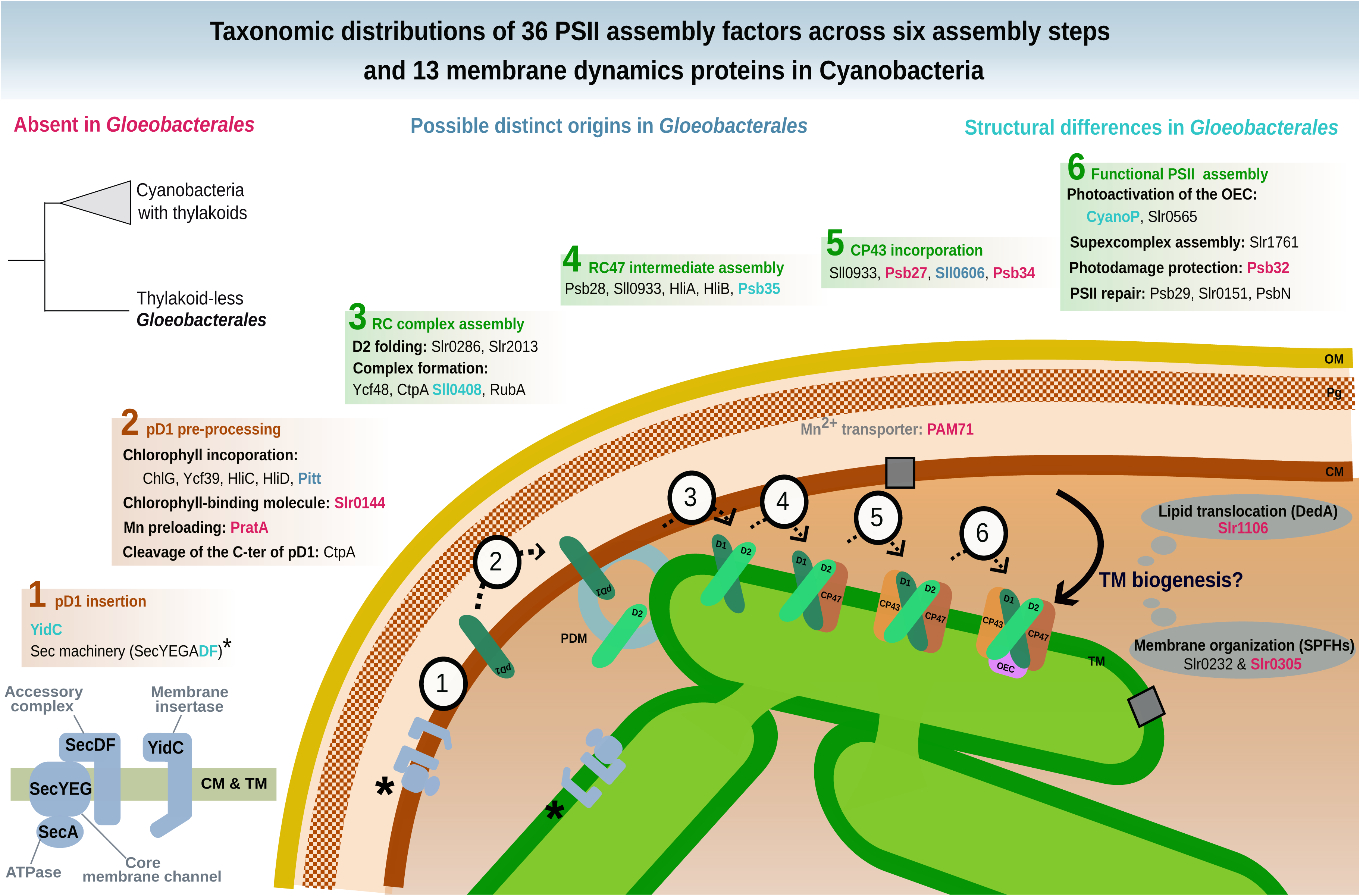
Overview of the results. Phylogenetic analyses of 36 assembly factors if PSII and 13 proteins involved in membranes dynamics summarized in (36) were performed to investigate the taxonomic distributions and structural differences between the thylakoid-less Gloeobacterales and the Phycobacteria.

Our results show that four additional proteins, from prohibitin and DedA families, involved in membrane trafficking, might constitute an alternative to the VIPP1-based model for the emergence of TM. Slr1106 is absent in *Gloeobacterales* but present in Phycobacteria, including *Thermostichales*, although its distribution within the latter group is limited. Slr1106 was acquired through lateral gene transfer (LGT) to the ancestor of Phycobacteria, but the donor could not be unambiguously determined (**Figures S144-S147**), possibly due to the ancient nature of the transfer. Nevertheless, alternative explanations, such as differential gene loss across cyanobacterial lineages or accelerated sequence evolution, cannot be completely excluded, albeit our species sampling is already very extensive. Slr1106 belongs to the prohibitin family, within the SPFH protein superfamily, typically associated with the inner mitochondrial membrane or the nucleus in eukaryotic cells, where they contribute to structural stabilization, apoptosis regulation, and cell signaling^55^. Members of the SPFH family are characterized by four α-helices and a twisted β-sheet structure, and they assemble into large oligomeric complexes^56^. Slr1106 is localized in both CM and TM^38^, consistent with a potential role in membrane organization and the formation of lipid microdomains. Besides Slr1106, the OG corresponding to the prohibitin Slr1106 includes proteins that are restricted to basal cyanobacteria, including *Gloeobacterales*, suggesting that these ancestral variants may have participated in the formation of primitive photosynthetic regions in CM, as observed in *Gloeobacterales*^5^. Another member of this family, Slr1768, was also identified within the same orthologous group and has already been shown to play a role in TM maintenance under high light conditions^57^. Slr1768 appears to be absent in *Gloeobacterales*, *Thermostichales* and *Gloeomargaritales* and only partially conserved across other cyanobacterial lineages **(Figure 3)**. In addition to these prohibitins, our analyses identified two DedA proteins that might have played a role in TM emergence: Slr0305 **(Figure 4)**, acquired at the root of Phycobacteria, and Slr0232 **(Figure 5)**, which exhibits an unusually long phycobacterial branch in its phylogeny. The DedA family contains three members in *Synechocystis* (Slr0232, Slr0305, and Slr0509). While this family is generally associated with membrane trafficking in eukaryotes, Slr0305 shows the highest homology to Tvp38, a putative lipid scramblase in yeast. Furthermore, Slr0305 is closely related to the *Arabidopsis thaliana* plastid protein AT5G19070.1, which is currently annotated as a SNARE-associated Golgi protein^58^. The phylogenetic distribution of these membranes trafficking proteins (emerging with *Thermostichales*), together with their reported function and membrane localization (known for Slr1106), is consistent with a role in TM emergence. This hypothesis contrasts with the CM–TM fusion model proposed for the ESCRT-III homolog VIPP1, whose central role in TM formation has been challenged by recent studies ^11,12^. However, our conclusions are based on comparative genomics analyses, and direct functional evidence is currently lacking for these candidate genes. Future experimental validation will be essential to clarify their precise roles and to determine whether they operate independently of, or inversely in coordination with, VIPP1.

The differences observed in late-stage PSII assembly factors are consistent with the respective membrane architecture of *Gloeobacterales* and Phycobacteria. Yet, evolutionary changes in assembly factors during cyanobacterial diversification, notably in photoprotection or PSII repair, could also have contributed to the patterns observed here, without necessarily being directly linked to thylakoid membrane emergence. On one hand, the absence (or distant *Gloeobacterales* homologs) of assembly factors related to the late stages of PSII biogenesis, namely Psb27, Psb34, Sll0606, may indicate an increased reliance on late-stage assembly steps during PSII repair in TM^59^. On the other hand, the absence of TM facilitates the biogenesis of the LET components, as it occurs within a single cellular compartment, the CM. This is particularly clear in the case of Mn utilization. Mn is stored in the periplasmic space^60^. In *Gloeobacterales*, the Mn cluster of PSII is located at the periplasmic side, and therefore does not require Mn transport across membranes for PSII assembly, unlike in Phycobacteria. The absence of PratA and PAM71 can thus be explained by the lack of TM. Interestingly, the SynPAM71 deletion mutant in *Synechocystis* sp. PCC6803 leads to a massive up-regulation of the Mn-binding CyanoP, which suggests a role of CyanoP both in PSII assembly and Mn homeostasis^50^. Our results show that *Gloeobacterales* CyanoP exhibits a less negatively charged surface compared with its orthologs in Phycobacteria **(Figure 8)**, which would suggest a change in Mn homeostasis from *Gloeobacterales* to Phycobacteria. Nevertheless, the Mn-binding protein Mn cupin A (MncA), which is associated with Mn storage in the periplasmic space^52^, might also be expected in *Gloeobacterales*, although we were unable to confirm its occurrence. Little is known about the function of the V4R domain of Slr0144 and Slr1471 in *Synechocystis*. Their absence in the majority of *Gloeobacterales,* along with duplication events at the base of Phycobacteria and within Thermostichales **(Figure 6)**, underscores unresolved questions regarding their roles in PSII assembly as part of the photosynthetically-related *slr0144*–*slr0152* operon^61^.

No close homologs of PSII assembly factor proteins were found for most photosynthesis-specific proteins In the non-photosynthetic sister lineages of Cyanobacteria **(Table S8)**. while all assembly factors involved in early stages of PSII biogenesis are already present in *Gloeobacterales*. Therefore, gene gain or loss cannot account for the relocation of PSII at the root of Phycobacteria. In contrast, the presence of additional helix loops in the predicted periplasmic domains of the phycobacterial YidC and SecD from *Thermostichales* might explain the differential localization of PSII between *Gloeobacterales* and Phycobacteria **(Figure 7)**. In *Escherichia coli*, YidC directly interacts with Sec components, particularly SecDF, which both contain periplasmic domains^47,62^. Interestingly, our data indicate that SecD of Phycobacteria harbors two additional helix–loop insertions in its periplasmic domain **(Figure S141).** In contrast, phycobacterial SecF differed only by its cytoplasmic domain in *Thermostichales*, whereas an additional helix–loop region was observed in *Synechocystis* sp. **(Figure S142)**. However, these structural interpretations are not empirical evidence of functional impact and remain hypothetical. At this stage, the interaction with YidC cannot be confirmed, as it has primarily been reported in *E. coli*, whose periplasmic domains differ from those found in cyanobacteria, thereby limiting direct structural comparison **(Figure S141C)**. Moreover, SecD localization within TM has not yet been established. Deletion of 90% of the periplasmic domain of *E. coli* YidC does not impair inner membrane protein biogenesis or cell viability^62^, and functional complementation experiments indicate that this domain is not strictly required for protein translocation, as Oxa1, despite lacking this domain, can complement YidC function in *E. coli*^63^. Altogether, these observations suggest that the periplasmic domain of YidC has a yet-undefined role in the cell, despite its conservation across diderm bacteria^64^. Cyanobacteria possess only a single set of Sec genes that operates in both the cytoplasmic membrane (CM) and the thylakoid membrane (TM), raising questions about how translocated proteins are selectively targeted to these two compartments^65,66^. In contrast, the YidC translocase is specifically localized to the PDM and TM. Since the lumenal space of Phycobacteria is homologous to the periplasmic space of *Gloeobacterales*^67^, the additional helix loops and structural modifications observed in the periplasmic domain of phycobacterial YidC could potentially underlie changes in its functional properties and contribute to YidC’s affinity for TM.

In the evolutionary scenario for the origin of TM, proposed by Cornet (2025) a series of intermediate stages allow transitioning from an electron transport chain located in the CM, i.e., the ancestral state still observed in *Gloeobacterales*, to the fully functional TM found in Phycobacteria (Cornet, 2025). This hypothesis is notably based on the high level of coordination among assembly factors, which reveals significant differences in the biogenesis of photosynthetic complexes, particularly between the two photosystems, suggesting distinct relocation processes^6^. In this framework, the first evolutionary stage corresponds to the formation of a membrane compartment capable of hosting relocalized photosynthetic complexes. At the other end, the final stage corresponds to the establishment of the fully developed phycobacterial TM system as it is observed today, whereas intermediate stages are hypothesized to involve alternative electron flux pathways. Our results on membrane trafficking proteins suggest an alternative to the VIPP1-based model for the first evolutionary stage. Indeed, they indicate that the emergence of compartmentalization may have involved proteins other than VIPP1. Alternatively, this first step may have required the prior establishment of a broader membrane context, including membrane-associated proteins and specific lipids, leading to the formation of early TM-like structures through vesiculation processes. Those structures could have supported simplified photosynthetic processes such as alternative electron flows, hence providing an evolutionary edge to these intermediates^6^. The observation that most PSII assembly factors are present in both *Gloeobacterales* and *Thermostichales* suggests that they were also encoded in the last common ancestor of extant Cyanobacteria, consistent with a long evolutionary history of oxygenic photosynthesis predating the diversification of modern Cyanobacteria^68,69^. This implies that the establishment of fully functional TM, the final stage, was driven primarily by structural innovations, such as those observed in YidC and SecD, rather than by gene gain. Once established, TM could have led to increased oxygen release into the atmosphere and potentially contributing to the GOE, as already suggested by Guéguen & Maréchal (2022).

Our results on PSII, the photosynthetic complex with the best-characterized biogenesis, highlight the value of evolutionary analyses of assembly factors for understanding the emergence of TM and the relocation of photosynthetic complexes from the ancestral CM-localized state to modern TM. Extending this approach to other complexes (PSI, the cytochrome *b6f*, ATP synthase, NDH-1) would provide a framework for elucidating intermediate evolutionary stages in TM emergence. Nevertheless, the biogenesis of these other complexes remains poorly characterized, with only a limited number of assembly factors identified to date^7^. This underscores the need to search for additional genes involved in the biogenesis of photosynthetic complexes if one wants to fully reconstruct the evolutionary emergence and diversification of the cyanobacterial photosynthetic apparatus.

## Conclusion

Our comparative analyses of membrane trafficking proteins and PSII assembly factors highlight key molecular innovations associated with the emergence of TM in Phycobacteria. The acquisition of prohibitins and DedA-family proteins, such as Slr1106, possibly via lateral gene transfer, and DedA-family proteins like Slr0305, probably facilitated the organization of lipid microdomains and the development of membrane compartments, providing a structural basis for thylakoids. In contrast, *Gloeobacterales* retain a simplified system, lacking V4R-domain proteins, late-stage PSII assembly factors (Psb27, Psb34, Sll0606), and manganese transporters, consistent with PSII biogenesis occurring entirely within the cytoplasmic membrane. Early-stage assembly factors are surprisingly conserved, but structural modifications in periplasmic domains of YidC and SecD in Phycobacteria may contribute to selective targeting of photosynthetic complexes to TM. Overall, thylakoid evolution may have required gene acquisitions for compartment formation and structural innovations enabling PSII migration.

## Author contributions

LH performed all analyses and drew figures. LH, DB, and LC wrote the manuscript. All authors read and approved the final version of the manuscript.

## Acknowledgments

We thank Edi Sudianto for kindly providing an additional dataset of Cyanobacteriota genomes and the corresponding species tree. This work was supported by a research grant (PDR T.0018.24 OR-OX-PHOT-IN-CYN) from the Belgian National Fund for Scientific Research (F.R.S.-FNRS) to DB. LC is supported by a mandate from the Belgian National Fund for Scientific Research (F.R.S.-FNRS). LH is a FRIA grantee of the Fonds de la Recherche Scientifique – FNRS. Computational resources were provided by the Consortium des Équipements de Calcul Intensif (CÉCI), funded by the F.R.S.-FNRS under Grant No. 2.5020.11 and by the Walloon Region. Molecular graphics and analyses performed with UCSF ChimeraX, developed by the Resource for Biocomputing, Visualization, and Informatics at the University of California, San Francisco, with support from National Institutes of Health R01-GM129325 and the Office of Cyber Infrastructure and Computational Biology, National Institute of Allergy and Infectious Diseases. ChatGPT (OpenAI) was used as a support tool for grammatical correction and text refinement.

## Data Availability

Multiple sequence alignments, phylogenetic trees, 3D structure predictions, and analysis scripts used for OrthoFinder analyses, orthologous group boxplots, phylogenetic trees, and ChimeraX are available on Figshare (DOI: https://doi.org/10.6084/m9.figshare.31135354). Detailed datasets of selected genome accessions are provided in Supplementary Table S4 (XLS format) and phylogenetic trees, including genome and protein accession numbers of orthologous proteins within Terrabacteria, are provided in Figs. S51–S95, together with homologous sequences within Cyanobacteriota in phylogenetic trees Figs. S96–S136. Table S4 summarizes the phylogenetic trees in the Terrabacterial and Cyanobacterial datasets from each orthologous groups of interests. The protein accession numbers supporting the findings of this study are, for CyanoP: BAC89381.1 (*Gloeobacter violaceus* PCC7421 GCA 000011385.1), MEN9209059.1 (Thermostichales cyanobacterium GCA_039794945.1) and the 3D PDB structure 2XB3 (*Thermosynechococcus vestitus* BP-1); and for YidC: BAC90172.1, MEN9209976.1 (Thermostichales cyanobacterium GCA_039794945.1), P74155.1 (*Synechocystis* sp. PCC 6803 substr. Kazusa). All other supporting data, including the Supplementary Figures file (containing phylogenetic trees, orthologous group selection and the ribosomal species tree) and the Supplementary Tables file (containing formatted tables and descriptions), are available in the Supplementary Information.

---

**Fig. S1** Phylogenomic tree based on ribosomal proteins.

**Fig. S2** Genome completeness by Terrabacteria phylum using CheckM2.

**Fig. S3** Genome contamination by Terrabacteria phylum using CheckM2.

**Fig. S4** Diversity curve generated using PARNAS v0.1.6 for the pruning of the ribosomal tree.

**Figs. S5–S50** Boxplots based on the bitscore distributions of BLASTp results.

**Figs. S51-S95** Gene trees of each OG of interest.

**Figs. S96-S136** Gene trees based on the best hits of a HMM search on the Cyanobacterial dataset (343 Cyanobacteriota species).

**Fig. S137** Additional β-sheet positioned outside of the ring structure formed during oligomerization in the proteins of the basal group I.

**Fig. S138** Interpro predictions of functional domains within the closest homologs of Sll0606 in Gloeobacterales.

**Fig. S139** Alignment of the cyanobacterial Slr0144 and Slr0147 sequences.

**Fig. S140** Alignment of the cyanobacterial YidC sequences.

**Fig. S141** AlphaFold3 predicted 3D structures of SecD from *Gloeobacter violaceus*, Thermostichales cyanobacterium GCA_039794545.1, *Synechocystis* sp. PCC 6803.

**Fig. S142** AlphaFold3 predicted 3D structures of SecF from *Gloeobacter violaceus*, Thermostichales cyanobacterium GCA_039794545.1, and *Synechocystis* sp. PCC 6803.

**Fig. S143** Alignment of the cyanobacterial CyanoP sequences.

**Fig. S144** Visualization of the selection of the sequences similar to Slr1106 with Ompa-Pa.

**Fig. S145** Visualization of the selection of the sequences similar to Slr1768 with Ompa-Pa.

**Fig. S146** Preliminary multigenic family tree of Slr1106.

**Fig. S147** Preliminary multigenic family tree of Slr1768.

**Table S1** Manual selection of the Terrabacteria representative species recovered from GTDB r220 (Parks et al., 2022).

**Table S2** Selection of the 343 genomes of Cyanobacteriota (Cyanobacterial dataset).

**Table S3** Sources of proteomes for the manually-selected accessions.

**Table S4** Selection of the 950 genomes based on a manual filtering and ribosomal tree pruning using PARNAS.

**Table S5** Description of selected assembly factors for this study.

**Table S6** Description of selected proteins involved in cyanobacterial membrane dynamics.

**Table S7** Summary of the BLASTp search and the phylogenetic trees.

**Table S8** Identification of homologs in closely related taxa.

