## Supplementary Figures for "Exploring Thylakoid Emergence: Evolution of Membrane Biogenesis and Photosystem II assembly in early-diverging Cyanobacteria"

**New Phytologist Supporting Information**
**Article title:** Exploring Thylakoid Emergence: Evolution of Membrane Biogenesis and Photosystem II assembly in early-diverging Cyanobacteria
**Authors:** Louise Hambücken, Denis Baurain, Luc Cornet
**Article acceptance date:** 27 April 2026

### Supplementary File

### Supplementary Figures

#### Genome selection and phylogenomic species tree

**Supplemental Figure S1: Phylogenomic tree based on ribosomal proteins. (A)** Uncollapsed tree and **(B)** collapsed tree by Terrabacteria phyla. Only bootstrap values <100 are shown. The tree was reconstructed from 1867 sequences across 3424 unambiguously aligned positions using IQ-TREE v2.4.0(Minh et al., 2020) with the best-fit substitution model (Q.YEAST+I+R10) and 1,000 ultrafast bootstrap replicates.

**Supplemental Figure S2: Genome completeness by Terrabacteria phylum using CheckM2(Chklovski et al., 2023).** The boxplot was created using the R package ggplot2 v4.0.0(Wickham et al., 2016) on Rstudio (2025.05.1 Build 513)(Posit team, 2025).

**Supplemental Figure S3: Genome contamination by Terrabacteria phylum using CheckM2.** The boxplot was created using the R package ggplot2 v4.0.0(Wickham et al., 2016) on Rstudio (2025.05.1 Build 513)(Posit team, 2025).

**Supplemental Figure S4: Diversity curve generated using PARNAS v0.1.6(Markin et al., 2023) for the pruning of the ribosomal tree.** The curve illustrates the relationship between the number of representative genomes retained and the phylogenetic diversity captured. Based on this analysis, a subset of 950 representative genomes was selected to maximize diversity while reducing redundancy in the dataset.

#### Identification of the orthologous groups of interest

**Figs. S5–S50: Boxplots based on the bitscore distributions of BLASTp(Camacho et al., 2009) hits for each (A) orthologous groups (OG) (B) hierarchical orthologous groups (HOG) from the N0 node inferred.** The OGs and HOGs were inferred using OrthoFinder v2.5.5(Emms & Kelly, 2019). The central line represents the median, the box bounds the interquartile range (IQR), whiskers extend to 1.5× IQR, and points beyond this range are shown as outliers. Boxplots were generated using the ggplot2 package in R (Wickham et al., 2016)

#### Orthologous gene tree inference

**Supplemental Figs. S51-S95: Gene trees of each OG of interest generated using IQ-TREE v3.0.0**[7] with the best-fit substitution model and 1,000 ultrafast bootstrap replicates. Bitscores were calculated by performing a BLASTp searches (v2.9.0+)(Camacho et al., 2009) of every protein from the 950 proteomes against a database containing reference sequences of known PSII assembly factors and proteins involved in membrane dynamics (**Table S8**).

**Supplemental Figs. S96-S136: Gene trees based on the best hits of a HMM search on the Cyanobacterial dataset (343 Cyanobacteriota species) with a HMM profile of the orthologous sequences of the proteins of interest.** The HMM search was performed using HMMER v3.4(Finn et al., 2011). The results of the HMM search was filtered using OmpaPa (https://metacpan.org/release/Bio-MUST-Core;D. Baurain).

**Supplemental Fig. S137: Additional β-sheet positioned outside of the ring structure formed during oligomerization in the proteins of the basal group I and II.** (A) AlphaFold3 predicted 3D structures of the SPFH proteins of the basal group I (BAC89785.1) and basal group II (BAC89786.1) from *Gloeobacter violaceus* with pLDDT scores. (B) AlphaFold3 predicted 3D structures of SPFH proteins from *Gloeobacter violaceus* (basal group I [BAC89785.1], blue, and basal group II [BAC89786.1], green) and two known SPFH proteins present in the same orthogroup from *Synechocystis* sp. PCC 6803 (Slr1106 [P72754], yellow, and Slr1768 [P73049], pink). (C,D) AlphaFold3 predicted 3D structures of the decamerization of BAC89785.1 with pLDDT scores. (E) Position of the extra β-sheet in the decamerization of BAC89785.1. The Predicted Local Distance Difference Test (pLDDT) scores mapped indicate confidence levels of the predicted models.

**Supplemental Fig. S138: Interpro Scan predictions(Jones et al., 2014) of functional domains within the closest homologs of Sll0606 in *Gloeobacterales*.** Functional protein domains were annotated with InterPro v5.48-83.0. The figure summarizes the domain architecture of each homolog.

**Supplemental Figure S139: Alignment of the cyanobacterial Slr0144 and Slr0147 sequences.** The alignment was performed using MAFFT v7.471 (Katoh, 2002). Residues that are 100% conserved are indicated by an asterisk (*); residues conserved in more than 50% of the sequences are indicated by a colon (:); and poorly conserved residues are indicated by a dot (·).

**Supplemental Figure S140: Alignment of the cyanobacterial YidC sequences.** The alignment was performed using MAFFT v7.471 (Katoh, 2002). The periplasmic domain as well as the extra alpha helices are indicated with braces. Residues that are 100% conserved are indicated by an asterisk (*); residues conserved in more than 50% of the sequences are indicated by a colon (:); and poorly conserved residues are indicated by a dot (·).

**Supplemental Figure S141: (A)** AlphaFold3 predicted 3D structures of SecD from *Gloeobacter violaceus* (GlSecD [BAC92107.1], blue), Thermostichales cyanobacterium GCA_039794545.1 (TSecD, yellow), *Synechocystis* sp. PCC 6803 (SynSecD, pink). **(B)** Additional helices identified in the periplasmic domain of SynSecD (pink). TSecD and GlSecD are shown respectively in blue and yellow. The Predicted Local Distance Difference Test (pLDDT) scores mapped onto the GlSecD, TsecD and SynSecD indicate confidence levels of the predicted models.

**Supplemental Figure S142: (A)** AlphaFold3 predicted 3D structures of SecF from *Gloeobacter violaceus* (GlSecF [BAC92107.1], blue), Thermostichales cyanobacterium GCA_039794545.1 (TSecF, yellow) cyan, and *Synechocystis* sp. PCC 6803 (SynSecF, pink). **(B)** Additional helices identified in the periplasmic domain of SecF from SynSecF. The Predicted Local Distance Difference Test (pLDDT) scores mapped onto the GlSecF, TsecF and SynSecF indicate confidence levels of the predicted models.

**Supplemental Figure S143: Alignment of the cyanobacterial CyanoP sequences.** The alignment was performed using MAFFT v7.471 (Katoh, 2002). Residues that are 100% conserved are indicated by an asterisk (*); residues conserved in more than 50% of the sequences are indicated by a colon (:); and poorly conserved residues are indicated by a dot (·).

**Supplemental Figure S144: Visualization of the selection of the sequences similar to Slr1106 with Ompa-Pa.** Matching sequences were plotted based on their sequence length in function of their -log10(e-value). The blue box corresponds to the selection (Copy per organism: 1 to 10, range of the fraction covering the HMM profile: 0 to 1, range of -log(evalue): 9 to 176, range of sequence length: 140 to 423). Sequences were visualized based on their (A) alignment coverage and (B) number of copies per genome.

**Supplemental Figure S145: Visualization of the selection of the sequences similar to Slr1768 with Ompa-Pa.** Matching sequences were plotted based on their sequence length in function of their -log10(e-value). The blue box corresponds to the selection (copy per organism: 1 to 10, range of the fraction covering the HMM profile: 0 to 1, range of -log(evalue): 13 to 175, range of sequence length: 149 to 456). Sequences were visualized based on their (A) alignment coverage and (B) number of copies per genome.

**Supplemental Figure S146: (A) Preliminary multigenic family tree.** The subtree corresponding to sequences similar to Slr1106 is shown in red. The tree was constructed from 11,714 sequences, corresponding to 319 unambiguously aligned positions, using FastTree(Price et al., 2010) with default parameters. The subtree highlighted represents the selected Slr1106 subtree, corresponding to 1380 sequences. **(B) Subtree limited to cyanobacterial sequences similar to Slr1106 (highlighted in red on panel A)).** The node support values are based on the Shimodaira-Hasegawa (SH) test (Gascuel, 1997).

**Supplemental Figure S147: (A) Preliminary multigenic family tree.** The subtree corresponding to sequences similar to Slr1768 is shown in red. The tree was constructed from 16,475 sequences, corresponding to 325 unambiguously aligned positions, using FastTree(Price et al., 2010) with default parameters. The subtree highlighted in blue represents the selected Slr1768 subtree, corresponding to 169 sequences. **(B) Subtree limited to cyanobacterial sequences similar to Slr1768 (highlighted in red on panel A)).** The node support values are based on the Shimodaira-Hasegawa (SH) test (Gascuel, 1997).s
