## Supplementary Tables for "Exploring Thylakoid Emergence: Evolution of Membrane Biogenesis and Photosystem II assembly in early-diverging Cyanobacteria"

**New Phytologist Supporting Information**
**Article title:** Exploring Thylakoid Emergence: Evolution of Membrane Biogenesis and Photosystem II assembly in early-diverging Cyanobacteria
**Authors:** Louise Hambücken, Denis Baurain, Luc Cornet
**Article acceptance date:** 27 April 2026

### Supplementary Tables

#### Genome selection and phylogenomic species tree

**Supplemental Table S1 (Excel sheets): Manual selection of the Terrabacteria representative species recovered from GTDB r220 (Parks et al., 2022).** Only the 43,827 genomes passing the GUNC contamination filter were retained. Within each GTDB order, genomes were ranked based on the difference between CheckM2 completeness and contamination scores. For orders represented by a single genus, the two highest-quality genomes were selected, whereas for all other orders, the top-ranked genome from each genus was retained. This procedure yielded a total of 1,452 genomes, excluding members of the phylum Cyanobacteriota.

**Supplemental Table S2 (Excel sheets): Selection of representative Cyanobacteriota genomes to complement the manual selection described in Table S1.** One or two genomes were selected per genus, prioritizing those with >80% completeness and <10% contamination, as estimated using CheckM2. When multiple genomes met these criteria, the highest-quality representatives were retained for downstream analyses.

**Supplemental Table S3 (Excel sheets): Sources of proteomes for the manually-selected accessions.** Proteomes were obtained either from NCBI RefSeq/GenBank annotations or predicted de novo using Prodigal v2.6.3 (Hyatt et al., 2010).

**Supplemental Table S4 (Excel sheets): Selection of 950 genomes based on manual filtering of Tables S1 and S2, combined with ribosomal phylogenetic tree pruning using PARNA (Markin et al., 2023).** A ribosomal phylogeny was used to guide genome subsampling, enabling the selection of a reduced, non-redundant dataset that maximizes phylogenetic diversity. The resulting set of 950 genomes was retained for downstream analyses.

#### Orthologous gene tree inference

**Supplemental Table S5 (This file): Description of selected assembly factors for this study.** Proteins were compiled from multiple literature sources and comprise factors implicated in Photosystem II (PSII) assembly, covering both early and late stages of the assembly pathway.

**Supplemental Table S6 (This file): Description of selected proteins involved in cyanobacterial membrane dynamics from (Siebenaller & Schneider, 2023).** Proteins were compiled from Siebenaller and Schneider (2023) and include proteins mplicated in membrane remodeling and organization.

**Supplemental Table S7 (This file): Summary of the BLASTp search and the phylogenetic trees.** Asterisks in the *Gloeobacter violaceus* PCC 7421 ortholog column means that this gene is present in *Gloeobacterales* but not in *Gloeobacter violaceus* PCC 7421.

**Supplemental Table S8 (This file): Identification of homologs in closely related taxa.** Identification of homologs in closely related taxa (Vampirovibrioniae, Sericytochromatia, and Margulisbacteria) based on HMM searches (Finn et al., 2011) performed across representative genomes from the GTDB (Parks et al., 2022). Highlighted proteins correspond to those for which no clear homologs were detected in any of these three lineages.

#### Supplemental Table S5.

| Category | Cyanobacterial gene of PSII assembly factors | Gene product name | Localization | Protein function | Target ligand or PSII subunits | Other interactions | Closest homolog in Arabidopsis thaliana | References |
|---|---|---|---|---|---|---|---|---|
| **precursor of the D19 pt protein (pD1) insertion** | slr1471 | YidC* | Intrinsic, PDM, TM | Membrane insertase | pD1 | Sec apparatus, ChlG-HliD complex, Ycf39, ribosome subunits | AT2G28800 (Alb3) | (Chidgey et al., 2014a; Gathmann, Sven et al., 2008; Proctor et al., 2018) |
| | sll1814 | SecY* | Intrinsic, CM, TM | Protein translocase subunit of the SecYEG channel | pD1 | Other proteins of the Sec apparatus, ribosome subunits, YidC | AT2G18710 | (Frain et al., 2016) |
| | ssl3335 | SecE | Intrinsic | Protein translocase subunit of the SecYEG channel | pD1 | | AT4G14870 | (Frain et al., 2016) |
| | NON20_19535 | SecG | Intrinsic | Protein translocase subunit of the SecYEG channel | pD1 | | - | (Frain et al., 2016) |
| | slr0774 | SecD | Intrinsic | Protein translocase subunit interacting with SecYEG channel | pD1 | | - | (Frain et al., 2016) |
| | slr0775 | SecF | Intrinsic | Protein translocase subunit interacting with SecYEG channel | pD1 | | - | (Frain et al., 2016) |
| | sll0616 | SecA | Intrinsic, CM (cytosolic side), TM (cytosolic side) | Protein translocase subunit with ATPase activity interacting with SecYEG channel | pD1 | | AT4G01800, AT1G21650 | (Frain et al., 2016) |
| **Chlorophyll-binding molecule** | slr0144 | Slr0144 | Intrinsic, TM | Putative chlorophyll-binding V4R domain | Chlorophyll (putative) | - | - | (Addlesee et al., 2000; Wegener et al., 2008) |
| **Chlorophyll incorporation into pD1** | slr0056 | ChlG* | Intrinsic | Terminal enzyme of the Chl biosynthesis pathway | Chlorophyll | Ycf39, HliD | AT3G51820 | (Proctor et al., 2018) |
| | slr0399 | Ycf39* | Intrinsic | Incorporation of the chlorophyll into pD1 | Chlorophyll precursors | HliD, pD1 | AT4G35250 (HCF244) | (Chidgey et al., 2014b; Linhartová et al., 2014; Sachelaru et al., 2013; Wysocka, 2025) |
| | ssl1633 | HliC | Intrinsic | Incorporation of the chlorophyll into pD1 | pD1 | HliD, Ycf39, YidC, ChlG, chlorophyll | CAB/HLIP/ELIP family | (Komenda & Sobotka, 2016) |
| | ssr1789 | HliD* | Intrinsic | Incorporation of the | Ycf39 | HliC, Ycf39, YidC, ChlG, | | (Konert, 2022) |

| | | | | | | | | |
|---|---|---|---|---|---|---|---|---|
| | | | | chlorophyll into pD1 | | chlorophyll, β-carotene | | |
| | slr1644 | Pitt | Intrinsic, PDM, TM | Tetratricopeptide Repeat Protein Involved in Light-Dependent Chlorophyll Biosynthesis | POR | - | AT1G78915 (TPR1) | (Schottkowski et al., 2009) |
| **Manganese-preloading interaction with pD1** | slr2048 | PratA* | Extrinsic, PDM, PP | Manganese pre-loading in pD1 | pD1, Mn2+ | CtpA | - | (Singh et al., 2004) |
| **Cleavage of the C-terminal extension of pD1** | slr0008 | CtpA* | Extrinsic, PDM | Peptidase cleaving the C-terminal extension of pD1 | pD1 | Associated with the PDM region | AT4G17740 | (Anbudurai et al., 1994; Karnauchov et al., 1997) |
| **D2 folding** | slr0286 | Slr0286 | Intrinsic, TM | Assembly and stability of the water-splitting system of photosystem II. | D2 | - | - | (Kufryk & Vermaas, 2001) |
| | slr2013 | Slr2013 | Intrinsic | Folding of the D2 protein | D2 | - | - | (Kufryk & Vermaas, 2003) |
| **RC complex formation** | slr2034 | Ycf48* | Extrinsic, L, PDM, TM | Stabilization of newly synthesized pD1, involved in PSII damage repair | pD1, iD1 | Sll0933 | AT5G23120 (HCF136) | (Kiss et al., 2019; Komenda et al., 2008; Rengstl et al., 2013) |
| | slr0008 | CtpA* | Extrinsic, PDM | Peptidase cleaving the C-terminal extension of iD1 | iD1 | Associated with the PDM region | AT4G17740 | (Anbudurai et al., 1994; Karnauchov et al., 1997) |
| | sll0408 | Sll0408 | Extrinsic, L | Peptidyl-prolyl cis-trans isomerase | CP47 | - | AT3G01480 (CYP38, TLP40) | (Fu et al., 2007; Sirpiö et al., 2008) |
| | D082_23770 | RubA | Intrinsic, TM | Rubdredoxin-line protein required for D1 and D2 association | D1, RC47-PSI | RC47-PSI complex | AT1G54500 (RBD1) | (Kiss et al., 2019; Šesták, 2007; Shen et al., 2002) |

| | | | | | | | | |
|---|---|---|---|---|---|---|---|---|
| **RC47 intermediate formation** | sll1398 | Psb28* | Intrinsic, CM, TM | Biogenesis of certain Chl-binding proteins | CP47 antennae, CP47 | - | AT4G28660 (PsbW) | (Bečková et al., 2017; Dobáková et al., 2009; Kashino et al., 2007; Sakata et al., 2013) |
| | sll0933 | Sll0933* | Intrinsic, TM | RC47 and PSII core formation | CP43, CP47 | Ycf48 | AT4G19100 (Pam68) | (Armbruster et al., 2010; Bučinská et al., 2018; Rengstl et al., 2011, 2013) |
| | ssl2542 | HliA | Intrinsic | Associated with later PSII assembly intermediate | CP47 antenna | HliC, chlorophyll, β-carotene | CAB/HLIP/ELIP family | (Akulinkina et al., 2015; Konert, 2022) |
| | ssr2595 | HliB | Intrinsic | | CP47 antenna | HliC, chlorophyll, β-carotene | - | (Akulinkina et al., 2015; Konert, 2022) |
| | ssl2148 | Psb35 | Putative intrinsic | Involved in in the life cycle of cyanobacterial Chl-binding proteins, especially CP47 | CP47 antenna, PSI trimer | - | - | (Pascual-Aznar et al., 2021) |
| **PSII lacking manganese cluster formation** | sll0933 | Sll0933* | Intrinsic, TM | RC47 and PSII core formation | CP47 antennae | Ycf48 | AT4G19100 (Pam68) | (Armbruster et al., 2010; Rengstl et al., 2011, 2013) |
| | slr1645 | Psb27* | Extrinsic, L | CP43 binding for assembly and Cluster oxygen-evolving complex assembly | CP43 antennae, D1 | - | AT1G03600 | (Komenda et al., 2012; Nowaczyk, 2014; Roose et al., 2007) |
| | sll0606 | Sll0606* | Intrinsic, CM or extrinsic, PP | Hypothetical transport of hydrophobic components;Insertion of the CP43 module into RC47 | CP43 antennae | - | - | (Komenda et al., 2008; Zhang et al., 2010) |
| | ssl1498 | Psb34 | Intrinsic | Lipoprotein involved in PSII late-assembly (competes with HliA/B for the same binding site) | CP47 assembled in PSII | - | - | (Rahimzadeh-Karvansara et al., 2022) |
| **photoactivation of the oxygen-evolving manganese cluster** | sll1418 | CyanoP (PsbP-like)* | Extrinsic, Lumen | Assembly of PSII intermediate, putative role in OEC assembly | D1, oxygen-evolving cluster | Divalent ions | AT3G55330 (PPL1) | (Knoppová et al., 2016; Nowaczyk, 2014) |
| | slr0565 | Slr0565 | Extrinsic, L | Catalyze of disulfide bond formation in lumenal and lumen-exposed proteins | PsbO | - | AT4G35760 (LTO1) | (Karamoko et al., 2011) |
| **Supercompl** | slr1761 | Slr1761 | Extrinsic, L | Fkbp-type Peptidyl- | PSII | Trx | AT3G60307 (FKBP-20-2) | (Lima et al., 2006) |

| | | | | | | | | |
|---|---|---|---|---|---|---|---|---|
| **ex assembly** | | | | prolylcis-transisomerase involved in PSII supercomplex assembly | | | | |
| **Protection from oxidative stress and photodamage** | sll1390 | Psb32 | Intrinsic, TM | PSII protection from photodamage and repair acceleration | PSII lacking manganese cluster | - | AT1G54780 (TLP18.3) | (Wegener et al., 2011) |
| **PSII repair** | sll1414 | Psb29 | Intrinsic, CM, TM | PSII repair (D1 degradation) | D1 ? | FtsH complex | AT2G20890 (THF1) | (Keren et al., 2005) |
| | slr0151 | Slr0151 | Intrinsic, TM | PSII repair under high-light conditions | CP43 antennae, D1 | - | - | (Yang et al., 2014) |
| | smr0009 | PsbN | Intrinsic, TM (stromal side) | Bitopic trans-membrane peptide required for PSII repair and PSII RC assembly | ? | ? | ATCG00700 | (Plöchinger et al., 2016; Torabi et al., 2014) |

#### Supplemental Table S6.

| **Category** | **Cyanobacterial gene of PSII assembly factors** | **Eukaryotic homolog** | **Protein function** |
|---|---|---|---|
| DedA proteins | Slr0232 | Tvp38 | Vesicular trafficking, putative lipid scramblase |
| | Slr0305 | | |
| | Sll0509 | | |
| SPFH proteins | Slr1106 | Prohibitin | Stabilization and dynamics of mitochondrial membrane, associated with lipid rafts |
| | Slr1768 | | |
| | Sll1021 | Flotilin | Associated with lipid rafts |
| | Slr1128 | Stomatin | Ion channel regulation, associated with lipid rafts |
| | Sll0815 | | |

| | | | |
|---|---|---|---|
| Cytoskeletal proteins | Sll1633 (FtsZ) | Tubulin | Microtubule formation |
| Membrane-bending proteins | Slr0483 | Arabidopsis CURT1 | Induction of curvature in TM |
| Dynamin-like proteins | Slr0869 | Dynamin | Membrane fission/fusion |
| ESCRT-III proteins | Sll0617 (IM30/VIPP1) | ESCRT-III | Membrane fission |
| | Slr1188 (PspA) | | Part of a membrane stress response system |

#### Supplemental Table S7.

| Gene name | Protein accession of the reference from NCBI | Orthogroup | *Gloeobacter violaceus* PCC 7421 (GCA_000011385.1) ortholog accession from NCBI | Bitscore | Phylogenetic tree of the corresponding orthogroup | Best-fit model for the orthogroup phylogenetic tree | Phylogenetic tree of the cyanobacterial sampling | Best-fit model fot eh phylogenetic tree of the cyanobacterial sampling |
|---|---|---|---|---|---|---|---|---|
| **Assembly factors of PSII** | | | | | | | | |
| YidC | P74155.1 | OG0000214 | BAC90172.1 | 342 | Fig. S94 | LG+F+R10 | Fig. S136 | Q.YEAST+I+R7 |
| SecY | P77964.1 | OG0000240 | BAC89334.1 | 527 | Fig. S71 | LG+F+R10 | Fig. S117 | LG+F+I+R7 |
| SecE | BAA17421.1 | OG0000328 | BAC88770.1 | 57.8 | Fig. S68 | Q.PFAM+R8 | Fig. S114 | Q.INSECT+F+I+R5 |
| SecG | BAA18612.1 | OG0000335 | BAC88772.1 | 73.6 | Fig. S70 | LG+F+R8 | Fig. S116 | Q.YEAST+F+R6 |
| SecD | BAA10118.1 | OG0000360 | BAC92107.1 | 458 | Fig. S67 | LG+F+R10 | Fig. S113 | LG+F+I+R8 |
| SecF | BAA10119.1 | OG0000411 | BAC92108.1 | 244 | Fig. S69 | LG+F+R10 | Fig. S115 | LG+F+I+R9 |
| SecA | BAA10347.1 | OG0000154 | BAC89777.1 | 1229 | Fig. S95 | LG+R10 | Fig. S112 | Q.YEAST+R10 |
| Slr0144 | BAA18544.1 | OG0003484 | - | - | Fig. S77 | LG+I+R5 | Fig. S138 | Q.PFAM+R6 |
| ChlG | QWO79731.1 | OG0002917 | BAC89750.1 | 421 | Fig. S51 | LG+F+R5 | Fig. S96 | Q.PFAM+I+R7 |
| Ycf39 | P74429.1 | OG0001181 | BAC91198.1 | 214 | Fig. S91 | Q.PFAM+F+I+R9 | Fig. S134 | LG+I+R7 |
| HliC | P73563.2 | OG0001968,OG00008795,OG0021297 | Unidentified | - | Fig. S55 | Q.PFAM+R7 | - | - |
| HliD | P72932.1 | OG0001968 | Unidentified | - | Fig. S55 | Q.PFAM+R7 | - | - |
| Pitt | BAA18461.1 | OG0003587 | BAC92203.1 | 61.6 | Fig. S57 | LG+F+I+R4 | Fig. S102 | LG+F+R7 |
| PratA | BAA17116.1 | OG0005899 | - | - | Fig. S58 | LG+F+R3 | Fig. S103 | LG+F+R6 |
| CtpA | Q55669.1 | OG0000126 | BAC88008.1 | 363 | Fig. S52 | Q.PFAM+F+R10 | Fig. S97 | LG+F+R10 |

| | | | | | | | | |
|---|---|---|---|---|---|---|---|---|
| Slr0286 | BAA18506.1 | OG0013412 | - | - | Fig. S80 | Q.PFAM+G4 | Fig. S124 | JTT+F+I+G4 |
| Slr2013 | BAA17260.1 | OG0001053 | -* | - | Fig. S89 | Q.PFAM+F+I+R9 | Fig . S137 | Q.PFAM+F+I+R7 |
| Ycf48 | BAA17091.1 | OG0004644 | BAC88796.1 | 218 | Fig. S92 | Q.PFAM+G4 | Fig. S135 | Q.YEAST+I+R7 |
| Sll0408 | AA10250.1 | OG0000241 | BAC89441.1 | 194 | Fig. S72 | WAG+R8 | Fig. S118 | Q.PFAM+I+R7 |
| RubA | P73068.1 | OG0001187 | BAC88795.1 | 100 | Fig. S66 | WAG+I+R5 | Fig. S111 | Q.PFAM+I+R5 |
| Psb28 | Q55356.3 | OG0005257 | BAC88869.1 | 56.6 | Fig. S60 | LG+G4 | Fig. S105 | Q.YEAST+I+R6 |
| Sll0933 | BAA16881.1 | OG0005529 | BAC89681.1 | 70.5 | Fig. S75 | Q.PFAM+F+I+G4 | Fig. S121 | Q.PFAM+F+I+R7 |
| HliA | P73183.1 | OG0001968 | Unidentified | - | Fig. S55 | Q.PFAM+R7 | - | - |
| HliB | P73429.1 | OG0001968,OG0009313 | Unidentified | - | Fig. S55 | Q.PFAM+R7 | - | - |
| Psb35 | BAA18333.1 | OG0006562 | BAC87990.1 | 47.4 | Fig. S64 | Q.PFAM+G4 | Fig. S109 | Q.PFAM+I+R5 |
| Psb27 | QWO79325.1 | OG0005894 | - | - | Fig. S59 | Q.PFAM+I+G4 | Fig. S104 | Q.PLANT+F+I+R6 |
| Sll0606 | BAA10364.1 | OG0000139 | (BAC90294.1) | 50.4 | Fig. S74 | Q.PFAM+F+R10 | Fig. S120 | |
| Psb34 | BAA18143.1 | OG0004508 | - | - | Fig. S63 | LG+G4 | Fig. S108 | Q.PFAM+I+R5 |
| CyanoP (PsbP-like)* | BAA18019.1 | OG0005391 | BAC89381.1 | 54.3 | Fig. S53 | Q.PFAM+G4 | Fig. S98 | Q.PFAM+R6 |
| Slr0565 | BAA10482.1 | OG0001631,OG0001293 | BAC90053.1 | 152 | Fig. S83 | WAG+F+R8 | Fig. S127 | LG+F+R7 |
| Psb32 | BAA18232.1 | OG00006067 | - | - | Fig. S62 | Q.PFAM+I+G4 | Fig. S107 | LG+F+R5 |
| Slr1761 | BAL34568.1 | OG0000856 | BAC88782.1 | 183 | Fig. S87 | WAG+F+R8 | Fig. S131 | WAG+F+I+R6 |
| Psb29 | BAA18023.1 | OG0005016 | BAC89341.1 | 164 | Fig. S61 | Q.PFAM+I+G | Fig. S106 | Q.PFAM+I+R7 |
| Slr0151 | BAA18551.1 | OG0001093 | - | - | Fig. S78 | LG+F+R8 | - | - |
| PsbN | P26286.1 | OG0005699 | BAC90942.1 | 50.1 | Fig. S65 | LG+I+G4 | Fig. S110 | MTZOA+I+G4 |
| PAM71 | P52876.1 | OG0001451 | - | - | Fig. S56 | LG+F+R8 | Fig. S101 | LG+F+R5 |
| **Proteins involved in membrane dynamics** | | | | | | | | |
| Slr0232 | BAA10237.1 | OG0000227 | BAC89450.1 | 137 | Fig. S79 | LG+F+R10 | Fig. S123 | Q.PLANT+F+I+R7 |
| Slr0305 | BAA10672.1 | OG0000999 | - | - | Fig. S81 | LG+F+R7 | Fig. S125 | Q.PLANT+F+I+R7 |
| Sll0509 | BAL36946.1 | OG0008882 | -* | - | Fig. S73 | Q.PFAM+I+G4 | Fig. S119 | Q.PFAM+F+I+R6 |
| Slr1106 | BFM39848.1 | OG0001215 | - | - | Fig. S85 | Q.PFAM+F+R8 | Fig. S129 | Q.PLANT+F+R6 |
| Slr1768 | BAA17070.1 | OG0001215 | - | - | Fig. S88 | Q.PFAM+F+R8 | Fig. S132 | Q.PLANT+F+G4 |
| Sll1021 | BAA16946.1 | OG0005308 | - | - | Fig. S76 | LG+F+G4 | Fig. S122 | Q.INSECT+F+R5 |
| Slr1128 | P72655.1 | OG0000457 | BAC90115.1 | 338 | Fig. S86 | LG+R10 | Fig. S130 | Q.PLANT+I+R7 |

| | | | | | | | | |
|---|---|---|---|---|---|---|---|---|
| Sll0815 | BAA18117.1 | OG0000457 | - | - | Fig. S86 | LG+R10 | - | - |
| Sll1633 (FtsZ) | P73456.1 | OG0000182 | BAC88239.1 | 469 | Fig. S54 | Q.YEAST+R10 | Fig. S99 | Q.INSECT+I+R9 |
| Slr0483 | BAA10316.1 | OG0005888 | - | - | Fig. S82 | LG+F+I+G4 | Fig. S126 | Q.PLANT+F+R7 |
| Slr0869 | BAA17817.1 | OG0010292 | - | - | Fig. S84 | LG+F+I+R3 | Fig. S128 | Q.PLANT+F+I+R7 |
| Sll0617 (IM30/VIPP1) | Q55707.1 | OG0001588 | BAC88839.1 | 217 | Fig. S90 | Q.YEAST+F+R7 | Fig. S100 | Q.INSECT+F+G4 |
| Slr1188 (PspA) | AGF53385.1 | OG0001588 | - | - | Fig. S90 | Q.YEAST+F+R7 | Fig. S100 | Q.INSECT+F+G4 |

#### Supplemental Table S8.

| **Protein** | **Vampirovibrioniae** | **Sericytochromatia** | **Marguslibacteria** |
|---|---|---|---|
| **PSII assembly factors** | | | |
| YidC | Yes | Yes | Yes |
| SecY | Yes | Yes | Yes |
| SecE | Yes | Yes | Yes |
| SecG | Yes | Yes | Yes |
| SecD | Yes | Yes | Yes |
| SecF | Yes | Yes | Yes |
| SecA | Yes | Yes | Yes |
| Slr0144 | No | Yes | No |
| **ChlG** | No | No | No |
| **Ycf39** | No | No | No |
| **Pitt** | No | No | No |
| PratA | Yes | Yes | No |
| **CtpA** | No | No | No |
| Slr2013 | Yes | No | No |
| **Slr0286** | No | No | No |
| **Ycf48** | No | No | No |
| **Sll0408** | No | No | No |
| **RubA** | No | No | No |
| **Psb28** | No | No | No |
| **Sll0933** | No | No | No |
| **Psb35** | No | No | No |
| **Psb27** | No | No | No |
| **Slll0606** | No | No | No |
| **Psb34** | No | No | No |
| Slr0565 | Yes | Yes | Yes |
| Slr1761 | Yes | Yes | Yes |

| | | | |
|---|---|---|---|
| **Psb32** | No | No | No |
| **Psb29** | No | No | No |
| **PsbN** | No | No | No |
| **Membrane dynamics** | | | |
| **Slr0509** | No | No | No |
| Slr0232 | No | Yes | No |
| Slr0305 | Yes | Yes | No |
| Sll1021 | No | Yes | No |
| **Slr1106** | No | No | No |
| **Slr1128** | No | No | No |
| **Slr1768** | No | No | No |
| FtsZ | Yes | Yes | Yes |
| **Slr0483** | No | No | No |
| **Slr0869** | No | No | No |
| **IM30 (VIPP1)** | No | No | No |
| PspA | Yes | Yes | No |
| **Manganese homeostasis** | | | |
| **CyanoP** | No | No | No |
| PAM71 | No | Yes | Yes |
